# Modeling Huntington’s disease against age-related genes reveals *CXXC4* as an epigenetic target to restore DRD1 striatal neuron activity

**DOI:** 10.64898/2025.12.12.693965

**Authors:** Maialen Arrieta-Lobo, Francesca Farina, Tamara Monteagudo Aboy, Megan Mair, Cloé Mendoza, Huy Tran, Jeff Aaronson, Jim Rosinski, Lisa Ellerby, Emmanuel Brouillet, Jean-Michel Peyrin, Juan Botas, Christian Neri, Lucile Megret

## Abstract

Neurons adapt gene expression to counter aging, yet the mechanisms by which they harness age-related genes to resist neurodegenerative disease remain elusive. We found that transcriptional-aging inversion (TAGI) in the *Drd1*-expressing striatal neurons (*Drd1* SNs) of Huntington’s disease (HD) knock-in (*Hdh*) mice exhibits discrete patterns against a TAG-like (TAGL) signature. Strikingly, TAGI dynamics may explain disease progression more accurately than TAGL. Moreover, in *Drd1* SNs, genes affected by 3’UTR accumulation during aging are predisposed to downregulation in aging and deregulation in *Hdh* mice. By integrating age-related 3’UTR data and *Hdh*-mice data, we identified a CAG-repeat-dependent network of upregulated genes with compensatory potential. This network features (i) *Atad-5,* a P CNA unloader that modifies CAG expansion in human HD plasma samples, and (ii) *CXXC4* (IDAX), an epigenetic regulator whose early-stage upregulation is lost as behavioral symptoms worsen in *Hdh* mice. Functionally, *CXXC4* reduces the senescence marker p16INK4a and restores glutamate excitability in human HD iPS cell-derived SNs. Collectively, these findings suggest that *Drd1* SN resilience capacity against HD relies on discrete age-related patterns and responses transiently activated in early disease stages. The reactivation of specific age-related genes such as *CXXC4* may notably restore cortico-striatal function in HD.

## INTRODUCTION

Neuronal vulnerability is a paradigm that is central to the study of neurodegenerative disease pathogenesis^1–4^. Besides vulnerability features that may be associated with specific neurons, vulnerability features may also be shared across vulnerable neuron populations^4^. Understanding the molecular underpinnings of neuronal vulnerability —particularly in disease-prone populations that are prone to repeat expansion diseases (REDs) such as striatal GABA-ergic neurons in Huntington’s disease (HD), a disease caused by CAG repeat expansion in huntingtin (Htt) that translates into polyglutamine (polyQ)-expanded Htt, primarily affecting the survival of striatal neurons^5^, has therefore strong therapeutic potential to sustain neuronal function, notably in the early phases of HD. Neurons may remodel gene expression in response to aging^6^, which might contribute to neuronal vulnerability or represent neuronal compensation. However, GABA-ergic neuron vulnerability over resiliency in the context of HD remains poorly understood on a systems level. Nonetheless, it appears that the CAG-repeat dependent loss of cellular resilience capacities^7^ and acceleration of molecular aging^8^ are two mechanisms that may drive HD in GABA-ergic neurons such as striatal neurons. Of note, aging-like effects of HD in neurons were previously suggested by advanced, however moderately, DNA-methylation age^9,10^ and by the acquisition of cellular senescence features in human HD medium-spiny neurons^11–13^. With regard to the cellular resilience capacities of neurons against HD, an important question is whether these capacities may actually involve genes that are altered by aging, *e.g.*, by reverting age-related expression patterns, and whether such a phenomenon may develop as a global response or series of discrete responses to mutant huntingtin. Another important question is about the critical time periods for such a cellular compensation mechanism to develop.

To address these questions, we performed a global analysis of the transcriptional changes entailed by HD on the genes altered by aging on transcriptional and 3’UTR accumulation levels in mouse *Drd1*-expressing striatal neurons (*Drd1* SNs). To this end, we developed a new computational approach that combines a network method based on spectral decomposition of the signal (SDS) for biological precision^14^ with bootstrappin for statistical robustness (Rando-SDS). We further combined Rando-SDS with a more powerful version of our shape-deformation analysis method *Geomic*^7^ for modeling the temporal dynamics of expression changes. Using these methods, we integrated information from (*i*) RNA-seq data from the allelic series of HD model knock-in (*Hdh*) mice —covering CAG-repeat-dependent TRAP-seq data obtained in *Drd1* SNs at 6 months of age^15^ and whole-striatum RNA-seq data across CAG repeat sizes and age points^16^— and (*ii*) age-related data, namely TRAP-seq data and 3’UTR accumulation data obtained in the *Drd1* SNs of older (24 months) wild-type mice^17^.

Our data on transcriptional aging-like and -inversion signatures in the *Drd1* SNs of *Hdh* mice suggest that the resiliency capacity of striatal neurons does not develop in the form of a global response that involves age-related genes at large, but rather develops in the form of discrete, early-stage and transient responses that involve age-related genes in a selective manner. Our data highlight the value of accounting for 3’UTR accumulation data in modeling the transcriptional effects of aging —and that of HD on age-related genes— in *Drd1* SNs. These findings noticeably suggest that restoring the expression of selected genes such as *CXXC4* (IDAX), an epigenetic regulator that is transiently upregulated in the *Drd1* SNs of *Hdh* mice and that is affected by 3’UTR accumulation in normal aging, may represent a promising therapeutic strategy to promote SN resiliency capacity against HD and perhaps other neurodegenerative diseases in which SNs are affected, *e.g*., spinocerebellar ataxias.

## RESULTS

### The *Drd1* striatal neurons of *Hdh* mice show discrete transcriptional aging-inversion patterns in the backdrop of aging-like gene-downregulation

To investigate the effects of HD on genes altered by aging in *Drd1* SNs, we compared ribosome-associated transcriptional changes that in these neurons are associated with normal aging^17^ and those associated increasing CAG repeat size in the allelic series of *Hdh* mice at 6 months of age^15^. We detected transcriptional aging-like (TAGL) effects including a significant overlap of 240 genes downregulated in both normal-aging and HD conditions and primarily annoted for synaptic transmission (**Figure 1a**, **Supplementary Table 1**), suggesting that HD and aging may converge to alter the functions of *Drd1* SNs via similar genes and pathways. In contrast, the TAGL-up component (158 genes) did not reach statistical significance, suggesting that these aging-like effects of HD are discrete and do not correspond to a global gene regulation mechanism. Transcriptional aging-inversion (TAGI) effects—whether of the up-to-down (230 genes up in aging and down in HD: synaptic plasticity) or down-to-up (95 genes down in aging and up in HD: cellular maintenance) type— did not show significant overlaps with aging-related transcriptional changes, suggesting that these aging-reverting effects of HD are discrete and do not correspond to a global regulation mechanism.

**Figure 1.**
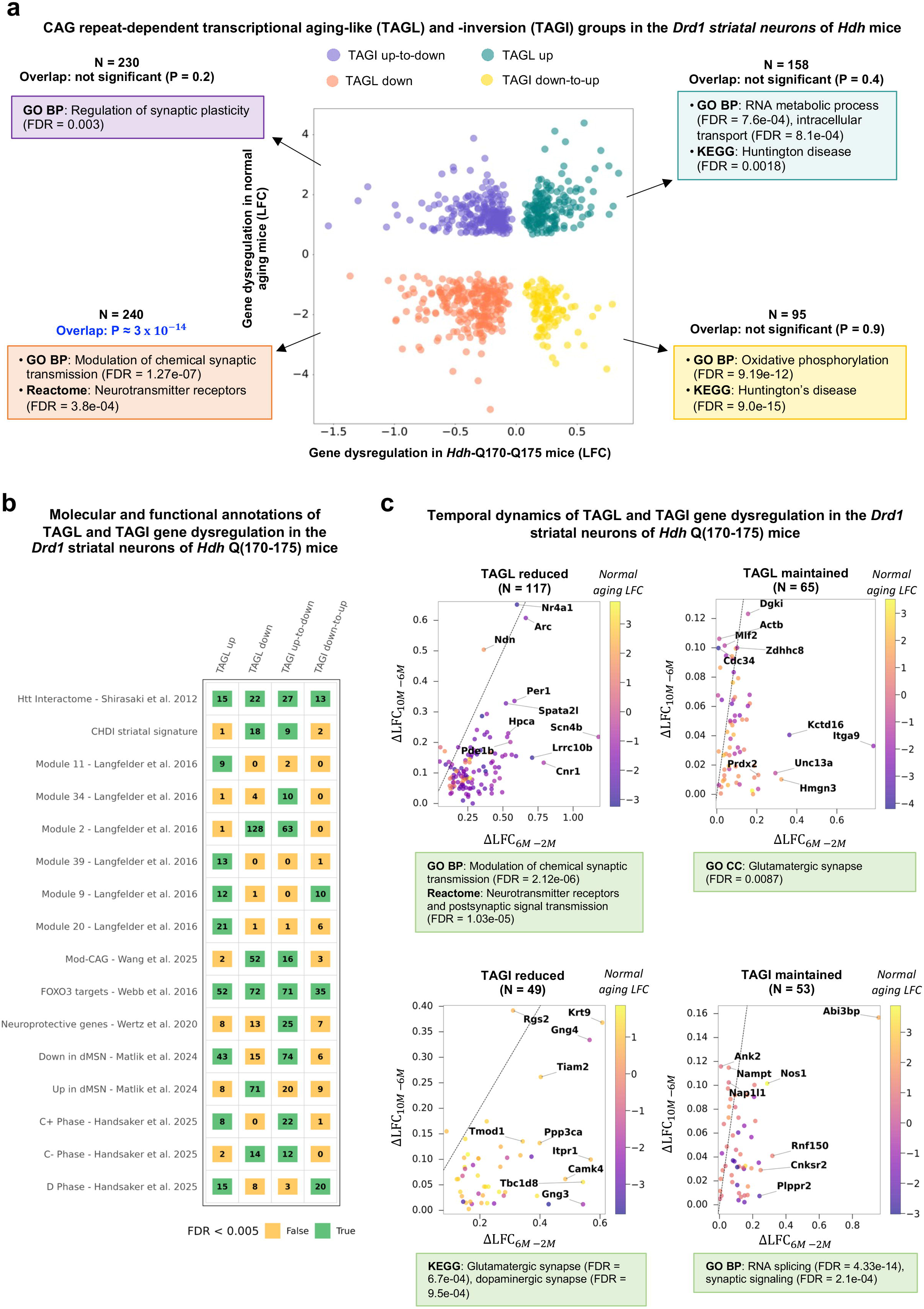
Characterization of CAG repeat-dependent transcriptional aging-like and - inversion components in the *Drd1*-expressing striatal neurons of *Hdh* mice. **(a)** Scatter plot showing log fold changes (LFC) in gene expression in the striatum of *Hdh* Q(170–Q175) mice (x-axis) versus naturally aged mice (y-axis). Each point represents a gene significantly dysregulated (FDR < 0.05) in both conditions. The number of genes and corresponding overlap P-values are indicated. P-values were computed by considering only genes that have a LFC, and thus were detected, in both aging and HD. The four resulting classes of tran-scriptional aging-like (TAGl) and -inversion (TAGI) groups are shown, including TAGl up (up regulation both in aging and HD), TAGL-down (down regulation both in aging and HD), TAGi-up-to-down (up regulation in aging and down regulation in HD) and TAGI-down-to-up (down regulation in aging and up regulation in HD). The most informative GO/BP and KEGG annotations (STRING database: https://string-db.org/) are indicated. (**b**) Overlap between TAGL-up, TAGI-up-to-down, TAGI-down-to-up and TAGL-down genes and transversal datasets of interest. The significances are represented with a color code, green if the Benjamini-Hochberg***-***corrected *P* values for multiple testing is lower than 0.005, red otherwise. The cardinal of the set is shown inside the corresponding square. (**c**) Temporal dynamics of TAGL and TAGI effects in the *Drd1*-expressing striatal neurons of Hdh mice. Out of 717 genes significantly dysregulated in the context of HD and that of normal aging (395 TAGL genes and 322 TAGI genes), 2239 genes (180 TAGl genes, 102 TAGi genes) were assigned at least to the *Drd1*-expressing striatal neurons using *Geomic* analysis. The panels show the four temporal subclasses defined by the difference between LFC values at 10-months *versus* 6 months (y-axis) compared to the difference between LFC values at 6-months *versus* 2 months (x-axis) The straight line in each panel represents the cancelation of effect at 10 months compared to 6 months. The most informative GO/BP and KEGG annotations (STRING database: https://string-db.org/) are indicated.

The four classes of transcriptional changes are enriched for HTT interactors^18^ (**Figure 1b**, **Supplementary Table 2**), suggesting that the transcriptional effects of HD on age-related genes involve genes and pathways proximal to huntingtin. They are also enriched in FOXO3 transcriptional targets^19^, a cell survival factor known to promote HD cell health^11^, suggesting that the transcriptional effects of HD on age-related genes may strongly involve genes that function in FOXO3 signaling, and highlighting FOXO3 transcriptional targets in the TAGL and TAGI classes as HD targets of special interest.

Additionally, TAGL and TAGI genes significantly overlap with several weighted gene coexpression network analysis (WGCNA) modules previously associated with CAG repeat length– and age-dependent transcriptional changes in the striatum of *Hdh* mice^16^ (**Figure 1b**, **Supplementary Table 2**). In particular, TAGL-down and TAGI up-to-down genes overlap with modules M2 and, to a lesser extent, M25—two downregulation modules related to neurotransmission, including cAMP signaling, postsynaptic density proteins, caudate markers, and glutamate receptor signaling. TAGI up-to-down genes specifically overlap with M34, a module enriched in transcription and chromatin factors. Conversely, TAGL-up genes overlap with M20 and M39, which are linked to stress responses and cellular homeostasis (M20: p53 signaling, cell division, protocadherins; M39: stress responses and DNA damage repair). Both TAGL-up and TAGI down-to-up genes are enriched in M9, a module associated with mitochondrial function, and TAGL-up genes specifically overlap with M11, a glia-related module downregulated in the bulk striatum of HD mice^20^, this latter overlap aligning with the idea that HD-associated transcriptional changes may display opposite patterns in neurons and glia. Of note, TAGL down and TAGI up-to-down genes are also enriched in ModCAG genes —a group of genes strongly altered by somatic expansion and largely downregulated in *Hdh* mice^21^ —, indicating that gene downregulation in the TAGL and TAGI classes may be associated with somatic expansion in the *Drd1* SNs of *Hdh* mice. These results suggest that the TAGL and TAGI classes are relevant to the molecular pathology of HD in the striatum, refining WGCNA module information. Interestingly, TAGI up-to-down genes are enriched for genes that promote striatal neuron survival in *Hdh-Q175* mice, but not for those that promote neuron death^22^, suggesting that TAGI up-to-down effects may promote the death of *Drd1* SNs in HD. Although the TAGI down-to-up class does not show enrichment in genes that promote striatal neuron survival in *Hdh-Q175* mice, This class does contain protective genes, raising the possibility that specific genes in the TAGI down-to-up class might protect from HD.

Finally, to assess the human disease relevance of TAGL and TAGI data, we compared these data to transcriptional and CAG repeat length data obtained from the striatal projection neurons of *post-mortem* human HD brains using fluorescence-activated nuclear sorting (FANS)^2^ or single-cell sequencing methods^23^. The TAGI down-to-up genes significantly overlap with genes downregulated in human HD direct striatal projection neurons (dSPNs)^2^, indicating opposite directions of dysregulation (**Figure 1b**, **Supplementary Table 2**). Interestingly, TAGL-up genes significantly overlap with genes upregulated in phase C, a phase associated with CAG expansions beyond 150 repeats in human HD dSPNs^23^. Additionally, TAGL-down genes significantly overlap with genes down-regulated in phase C, which also applies to TAGI-up-to-down genes^23^. Finally, TAGL up and TAGI down-to-up are enriched in phase D genes, where phase D is characterized by a transcriptional de-repression crisis associated with very large CAG repeats in human HD dSPNs^23^. Together, these results suggest that while the TAGL and TAGI signatures in *Hdh* mouse *Drd1* SNs are discrete (except to TAGL down), they are consistently (same direction of deregulation) relevant to several phases of the molecular pathology of human HD dSPNs.

### The *Drd1*-expressing striatal neurons of *Hdh* mice maintain transcriptional aging-inversion while aging-like changes are reduced

Next, we assessed the temporal dynamics of the TAGL and TAGI effects from 2 months to 6 months (weakly symptomatic mice) and 6 months to 10 months (strongly symptomatic mice) of age. To this end, we used our software *Geomic*, a shape deformation analysis method that infers information on the temporal dynamics of molecular changes via statistically matching the shapes of temporal data (*e.g*., available for whole-striatum gene expression surfaces) with that of CAG repeat-dependent data (*e.g*., cell type-specific gene expression curves)^16^. To increase gene coverage, we developed a more powerful version (v 1.1) of *Geomic* (see Methods). Using *Geomic* v 1.1, whole-striatum gene-expression surfaces (covering CAG-repeat lengths and age points in *Hdh* mice) could be matched to CAG repeat-dependent gene expression curves at six months of age and therefore assigned to *Drd1* SNs for a large number of TAGL and TAGI genes. Temporal information (dysregulation maintained or reduced) was retrieved for 45.7% (182/398) of TAGL genes and for 31.3% (102/325) of TAGI genes, all of them primarily associated with neurotransmission (**Supplementary Table 1**). The distribution of the maintained and reduced subclasses is significantly different between TAGL and TAGI (p-value = 0.011, chi2 test using R version 4.5.0). We found that the majority of the TAGL effects are reduced as *Hdh* mice become strongly symptomatic, which primarily applies to down-regulated genes such as for example *Per1* and *Scn4b* (**Figure 1c**, **Supplementary Table 1)**. In contrast, about half of the TAGI effects for which temporal information could be retrieved was maintained (primarily involving genes down-regulated in *Hdh* mice), with the other half that was reduced (primarily involving genes up-regulated in *Hdh* mice). These results suggest that, compared to TAGL dynamics, TAGI dynamics may better explain disease progression in the *Drd1* SNs of *Hdh* mice. Since the TAGI up-to-down class is enriched in genes that promote striatal neuron death when reduced in *Hdh* mice (see **Figure 1b**), these results also suggest that one effect of the maintenance of TAGI effects is to drive the disease in *Drd1* SNs.

### Genes downregulated in aging are more likely affected by 3’UTR accumulation in mouse *Drd1*-expressing striatal neurons

Besides transcriptional dysregulation, 3’UTR accumulation is another feature associated with the effect of normal aging on *Drd1* SNs^17^. Approximately 5,000 genes in *Drd1*-SNs of older mice were previously reported to undergo 3′UTR accumulation^17^, a phenomenon that can lead to reduced protein levels as illustrated for *Rras2* in senescent cells^24^. While data on the effects of HD on 3’UTR accumulation are not available, we reasoned that modeling the effects of HD on aging-related genes would gain to account for information available on 3’UTR accumulation in the course of aging in *Drd1* SNs. We first tested for a link between TRAP-seq data and 3’UTR accumulation data obtained from *Drd1* SNs in the context of normal aging. Interestingly, we found that genes downregulated (but not upregulated) in aging *Drd1* SNs are more likely affected by 3′UTR accumulation as indicated by a significant overlap of 173 genes enriched for genes associated with the mitochondrial respirasome (**Figure 2a**, **Supplementary Table 3**). Additionally, based on using Enrichr^25^, genes upregulated and affected by 3′UTR accumulation are enriched in transcription factors that are also HTT interactors, including NFATC1 and NR3C1 (Transcription Factor PPIs; FDR < 0.05). In contrast, downregulated genes affected by 3′UTR accumulation are associated with repressive histone marks such as H3K27me3 and H3K36me3 (ENCODE histone modification/kidney or mouse lymphoma cell lines CH12.LX, FDR < 0.01) and for transcriptional targets of SP1, a transcription factor that binds to HTT and that is strongly implicated in HD pathogenesis^26^. Together, these results suggest that, in aging *Drd1* SNs, downregulated genes are preferentially affected by 3′UTR accumulation, are likely under SP1 regulation, and could undergo epigenetic repression, highlighting a potential convergence of epigenetic, transcriptional and post-transcriptional effects during neuronal aging of striatal neurons, particularly for reduced expression.

**Figure 2.**
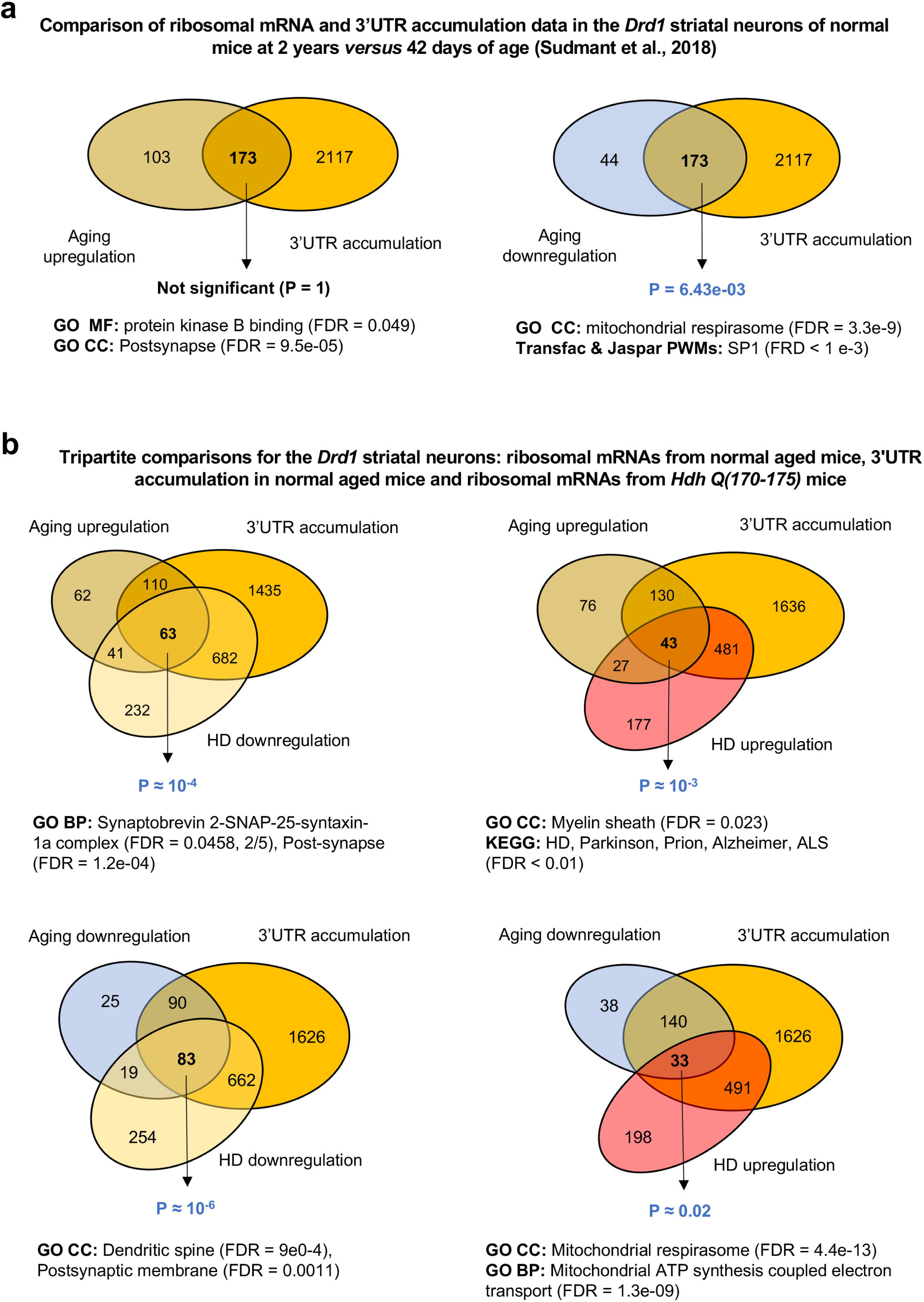
Overlaps between gene dysregulation in normal aging, 3’UTR accumulation in normal aging, and HD-associated gene dysregulation in mouse *Drd1*-expressing striatal neurons. (**a**) Overlaps between gene dysregulation and 3’UTR accumulation in normal aging^17^. Left panel: overlap between genes upregulated in normal aging (genes with adjusted P-values ≤ 0.05 were considered significant) and genes affected by 3’UTR accumulation during aging. Right panel: overlap between genes that are down regulated during normal aging (genes with adjusted P-values ≤ 0.05 were considered significant) and genes affected by 3 ’UTR accumulation during aging. (**b**) Three-way overlaps covering genes dysregulated (genes with adjusted P-values ≤ 0.05 were considered significant) in the *Drd1* SNs of *Hdh* mice^15^. The four categories (TAGl up, TAGI up-to-down, TAGI down-to-up, TAGL down) previously identified (see Figure 1) are significantly enriched in genes affected by 3’UTR accumulation in normal aging.

### Genes deregulated in the *Drd1*-expressing striatal neurons of *Hdh* mice are more likely dysregulated and affected by 3’UTR accumulation in normal aging

Detection of a significant overlap between gene dysregulation and 3′UTR accumulation in *Drd1* SNs during normal aging, —encompassing transcription factors implicated in HD (**Figure 2a**)—, led us to hypothesize that genes showing ribosome-bound mRNA dysregulation in *Hdh* mice substantially coincide with those dysregulated and affected by 3′UTR accumulation under normal aging conditions. Probabilistic overlap analysis indicated that this hypothesis is true for all four tripartite comparisons of these datasets (**Figure 2b**, **Supplementary Table 4**). This pattern was particularly significant for 83 TAGL-down genes associated with synaptic transmission, which might correspond to a detrimental aging-like effect of mutant huntingtin in these neurons. This pattern was also significant for 33 TAGI-down-to-up genes that are associated with the mitochondrial respirasome, which might correspond to a survival-promoting aging-inversion effect in these neurons. These results suggest that, in the *Drd1* SNs, genes dysregulated and affected by 3’UTR accumulation in aging are more likely deregulated in *Hdh* mice, and this phenomenon may apply to both TAGL and TAGI effects in these mice.

### Graph-based integration of gene dysregulation data in *Hdh* mice and 3’UTR accumulation data in aging defines biologically distinct networks

Given that genes deregulated in the *Drd1* SNs of *Hdh* mice are more likely those that are altered on transcriptional and post-transcriptional levels in normal aging and this may involve biological processes directly relevant to HD pathogenesis (**Figure 2b**), we investigated the relationships between genes deregulated in *Hdh* mice and genes affected by 3’UTR accumulation in aging on a systems level. To this end, we developed Rando-SDS, a network analysis work-flow that includes a bootstrapping step downstream to the spectral decomposition of the signal^14^ —here, ribosome-associated mRNA levels from *Hdh* mice at 6 months of age ^15^— against a probabilistic functional network (here, the STRING mouse network). The 3′UTR accumulation data for *Drd1* SNs in normal aging — represented by the R_tc_ scores where an R_tc_ of 1 represents a two-fold enrichment in 3′UTR versus CDS sequences^17^— were then projected onto these Rando-SDS networks (**Figure 3a**, **Supplementary Tables 5-7**; see also Methods).

The Rando-SDS analysis defined two interconnected networks for Q111 and for Q(170-175), including one network for genes downregulated in *Hdh* mice and affected by 3′UTR accumulation in aging (Class-1 network), and one network for genes upregulated in *Hdh* mice and affected by 3′UTR accumulation in aging (Class-2 network). The CS-1 score and CS-2 score account for the relative strength of gene dysregulation in HD *versus* 3′UTR accumulation in normal aging for each Class-1 and Class-2 gene, respectively (**Figure 3a, Supplementary Table 6**). The size of the Class-1 network is dependent on CAG-repeat lengths (Q50: 6 genes; Q111: 80; Q170: 184; Q175: 189, **Supplementary Table 7**). Top annotations for the Q(170-175) Class-1 genes (N = 246) are histone deacetylase complexes and neurotransmission processes, with high-scoring genes such as the small GTPase *Rhob* (CS-1 = 0.75) and the kinase *Dclk3* (CS-1 = 1.11), a striatal gene that protects neurons against HD^27^, several of which promote synaptic transmission such as for example the neuronal gene *Arc* (CS-1 = 1.03). Similarly to TAGL genes (**Figure 1b**), Q(170-175) Class-1 genes showed a strong, CAG-repeat-dependent overlap with WGCNA module M2 (Langfelder et al., 2016 PMID: 26863350, **Figure 3b, Supplementary Table 8**), which is associated with CAG repeat length and dysregulation of genes associated with neurotransmission (cAMP signaling, postsynaptic density proteins, caudate markers). The size of the Class-2 network is also dependent on CAG-repeat lengths (Q50: 19 genes; Q111: 74; Q170: 99; Q175: 115, **Supplementary Table 7**). Top annotations for the Q(170-175) Class-2 genes (N = 143) are endosomal and vesicular trafficking—notably endosomes, multivesicular bodies, and extracellular vesicles—with high-scoring genes that are strongly-deregulated — a rare event— such as the transcription factor *Sox11* (CS-2 = 0.68, LFC of about 0.6), and mid-scoring genes that are less strongly-deregulated —a more frequent and more representative event—, such as the epigenetic regulator *CXXC4* (CS-2 = 0.43, LFC in the range of 0.2-0.3), along with additional genes that may promote synapse formation (tyrosine phosphatase *Ptpro,* CS-2 = 0.57) and neuronal homeostasis (Fidgetin, CS-2 = 0.74). As observed for TAGI genes (**Figure 1b**), Q(170-175) Class-2 genes overlap with WGCNA modules M20 (p53 signaling, cell division, protocadherins) and M9 (mitochondrial function) (**Figure 3b, Supplementary Table 8**). Of note, the Class-2 network is enriched for targets of CTCF, a chromatin regulator that maintains genomic insulation and regulates gene expression in neurons through the control of 3D chromatin architecture and that may protect against CAG expansion in HD^28^. This enrichment raises the possibility that several Class-2 genes (upregulated genes) may reflect an adaptive compensatory mechanism mediated by CTCF.

**Figure 3.**
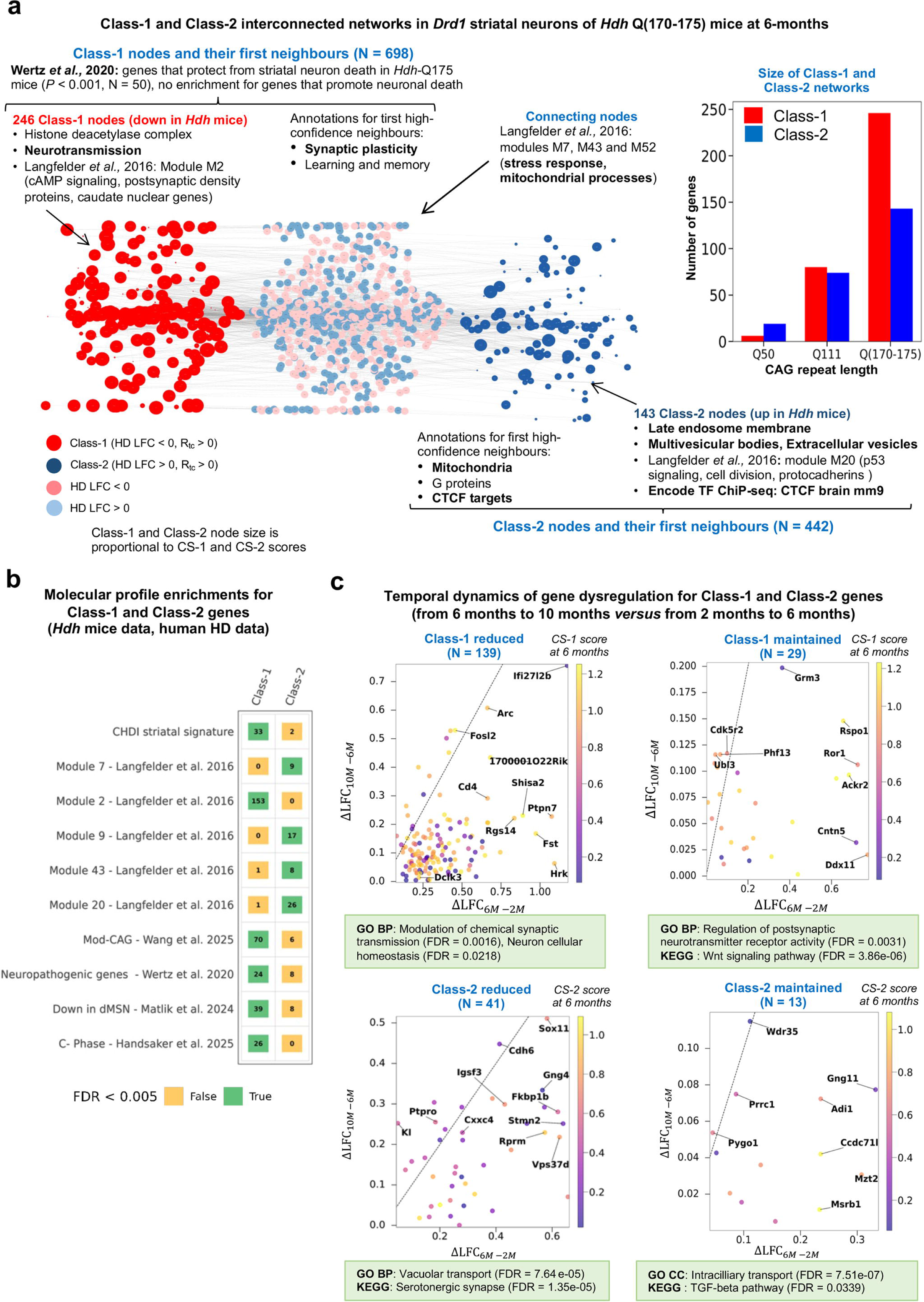
Graph-based integration of gene dysregulation data in *Hdh* mice and 3’UTR accumulation data in aging for mouse *Drd1*-expressing striatal neurons. (**a**) Interconnections between the Q(170-175) Class-1 (red) and Class-2 (green) gene nodes. Nodes downregulated in HD but not present in the Class-1 network are light red, while nodes downregulated in HD but not present in the Class-2 network are light green. Node size is proportional to the CS-1 or CS-2 scores for Class-1 or Class-2 gene nodes, respectively. The most informative EnrichR annotations are indicated, as well as the enrichment in WGNA modules^16^. (**b**) Overlaps between Class-1 or Class-2 genes and transversal datasets of interest. FDR values lower than 0.005 were considered significant. (**c**) Temporal dynamics of Class-1 and Class-2 gene expression in the *Drd1*-expressing neurons of *Hdh* Q(170-175) mice. Out of 389 genes retained by our model in the combined Q(170-175) network (246 Class-1 genes and 143 Class-2 genes), 222 genes (168 Class-1 genes and 54 Class-2 genes) were assigned at least to the *Drd1* SNs. These genes are classified into two temporal subclasses (educed or maintained) for Class-1 and Class-2 as defined by the difference between LFC values at 10-months *versus* 6 months (x-axis) compared to the difference between LFC values at 6-months *versus* 2 months (y-axis). The straight line in each panel represents the cancelation of deregulation between 6 months and 10 months compared to the deregulation between 6 months and 2 months. The most informative GO/BP and KEGG annotations (STRING database: https://string-db.org/) are indicated where 29 high-confidence STRING neighbors have been added to Class-1/’maintained’ genes, 49 high-confidence STRING neighbors have been added to Class-2/’reduced’ genes and 13 high-confidence STRING neighbors have been added to Class-2/’maintained’ genes.

Adding high-confidence neighbors (see Methods) in equal proportion compared to seed nodes (**Supplementary Table 7**) revealed that the Class-1 network is associated with synaptic plasticity, learning, and memory processes, whereas the Class-2 network is associated with mitochondrial and G-protein pathways (**Figure 3a**). Noticeably, the expanded Class-1 network is enriched in genes that protect mouse striatal neurons from the HD process (Wertz et al., 2020, **Supplementary Table 9**), suggesting that downregulation of Class-1 genes may contribute to pathogenicity. Similar to the Class-2 network, the expanded Class-2 network is enriched for targets of CTCF, further raising the possibility that upregulation of Class-2 genes may reflect an adaptive compensatory mechanism mediated by CTCF targets.

Finally, the Class-1 and Class-2 networks were interconnected through shared nodes enriched in WGCNA modules M7, M43, and M52, which relate to stress responses and mitochondrial processes. The shortest path length (SPL) between network nodes increased from SPL = 1 at Q50 to SPL = 4 in the combined Q(170-175) network, suggesting that the biological mechanisms in which Class-1 and Class-2 genes are involved may become increasingly distinct as CAG-repeat length increases in *Drd1* SNs.

Together, these results suggested that while Class-1 genes show biological features relevant to HD pathogenicity, Class-2 genes show biological features relevant to disease resistance.

### The majority of Class-1 and Class-2 transcriptional changes are reduced as *Hdh-Q(170-Q175)* mice become strongly symptomatic

To determine whether gene deregulation in the Class-1 and Class-2 components may be reduced or maintained as Hdh mice move from weakly (2-6 months of age) to strongly (6-10 months of age) symptomatic phases, we used an improved version (v 1.1) of the shape deformation analysis software *Geomic* (see Methods). The *Geomic* v 1.1 analysis enabled information about the temporal dynamics of gene dysregulation to be retrieved for 68.2% (168/246) of Q(170-175) Class-1 genes. Similarly, temporal information was recovered for 37.7% (54/143) of Q(170-175) Class-2 genes, This set of information was relevant to 42 Class-1 genes and 13 Class-2 genes that were also detected for the Q111 condition. The upregulation at 6 months was reduced at 10 months for the majority of Class-1 genes for which information on temporal changes of expression was obtained, and the same conclusion applied to Class-2 genes for which information on temporal changes of expression was obtained . Thus, both Class-1 and Class-2 genes are more likely to be reduced (chi2 non-significant). (**Figure 3c, Supplementary Table 6**). Assuming that Class-2 genes might primarily be involved in disease resistance whereas Class-1 genes may primarily contribute to pathobiology (**Supplementary Table 9**), these results raise the possibility that the reduction of Class-2 effects from 6 months to 10 months of age may better explain disease progression compared to the reduction of Class-1 effects. Along these lines, Class-2 genes of interest are those for which upregulation at 6 months of age is basically lost at 10 months of age, notably *Sox11* (SRY-Box Transcription Factor 11), the top CS-2 score gene and a gene that protects striatal neurons from cell death in wild-type mice^22^, and median CS-2 score yet strongly-deregulated genes such as *CxxC4*, an epigenetic regulator able to repress of TET2^29^, and PtprO, a tyrosine phosphatase associated with synapse formation (**Figure 3c**).

### Class-2 and Class-1 genes modify motor impairement in transgenic *HTT* flies

To assess whether genes in the Class-1 and Class-2 signatures can modulate HD pathology, we assessed whether the genetic manipulation of 80 selected (medium to high CS score) Class-1 and Class-2 genes could modulate neuronal dysfunction induced by mutant *HTT* (mHTT) a Drosophila melanogaster model^30,31^. Flies expressing an expanded N-terminal fragment of human *HTT* (*HTTNT231Q128*) exhibit motor impairments that progress longitudinally, which are quantifiable using the startle-induced negative geotaxis response (**Figure 4a**). Through this model, we can determine whether manipulation of the selected Class-1 and Class-2 genes worsens or ameliorates mHTT-induced neuronal deficits.

**Figure 4.**
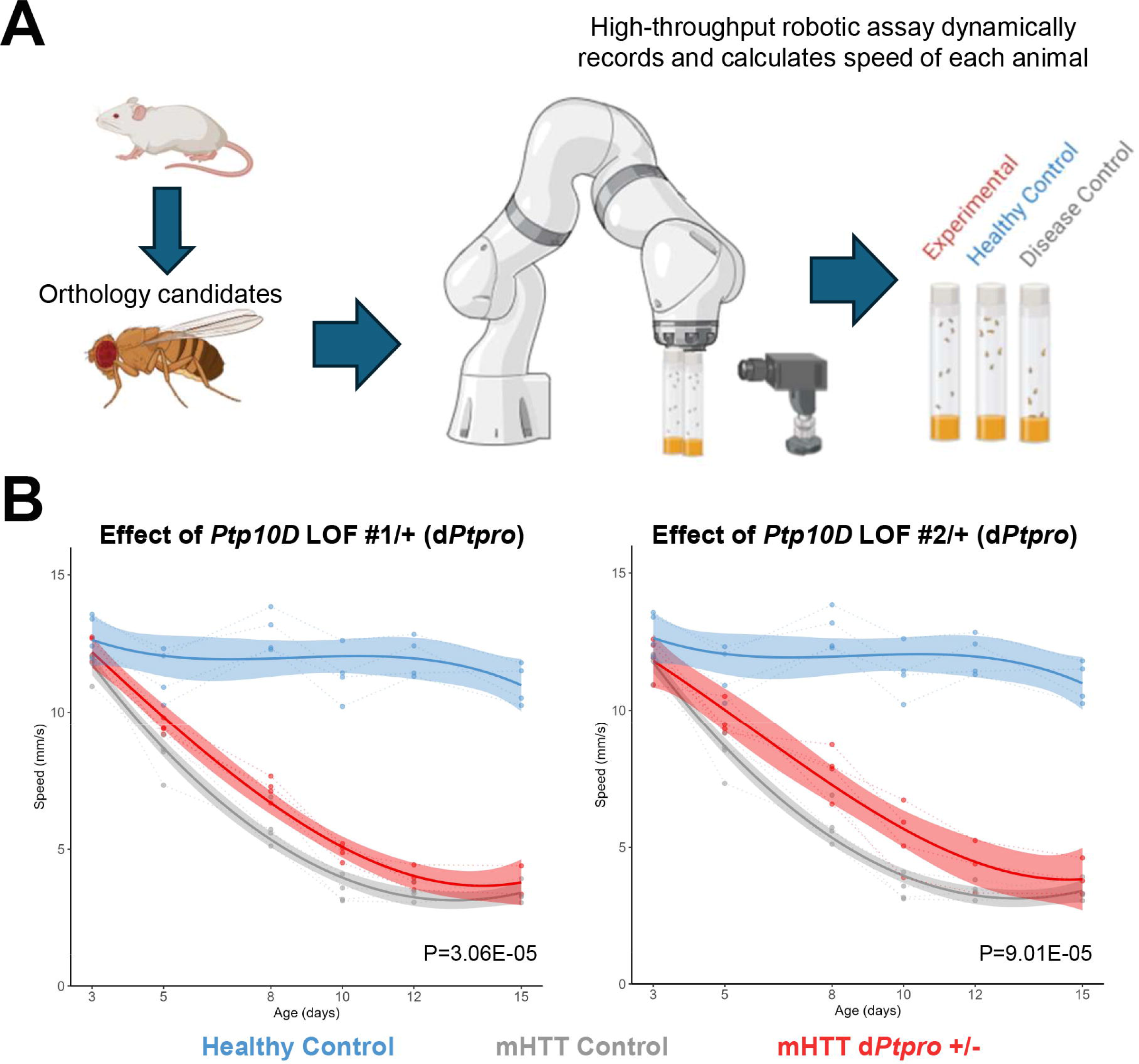
Network cluster connecting mouse Class-2 gene *Atad-5* and six genes human genes upregulated in the *Drd1* striatal projection neurons of human HD brains. Network cluster generated using six human genes encoding integral components of the PCNA complex and upregulated in the dSPNs of *post-mortem* human HD brains including *CDKN1A*, *EP300*, *DONSON*, *PRIM1*, *SHPRH* and *POLD1*^2^ and the human ortholog of Class-2 gene *Atad-5* as seed genes with addition of 50 1st neighbors distributed on 1^st^ shell or 2^nd^ shell in protein complexes as extracted from the STRING database^47^. Inserts show the CAG-repeat and time-dependent of pattern of expression (http://www.broca.inserm.fr/BrainC_database/) of *Atad-5* and that of two other genes of interest that in the *Drd1* SNs of *Hdh* mice that shows upregulation at 6 month of age followed by reduction of this increase at 10 months of age^7^ including (*i*) *CTF18*, a clamp loader bound to PCNA^36^ that promote touch receptor neuron dysfunction in human exon-1 *HTT C. elegans* nematodes^32^ and for which upregulation in the *Drd1* SNs of *Hdh* mice is pathogenic as indicated by the *Geomic* analysis^7^, and (*ii*) *Brd4*, a negative regulator of PCNA unloading^37^ which binds and inhibits the ATAD-5 complex^38^ and is however not referenced in the STRING database.

In total, 52 fly orthologs of 44 human genes were assessed for their potential to modulate mHTT-induced neuronal dysfunction. Across the two categories of genes, we identified 24 fly genes that modified the health of the animals (**Supplementary Table 10**). In the Class-1 group, we identified manipulation of two target genes worsened the neuronal dysfunction phenotype and two induced lethality, while loss-of-function of six candidate reduced neuronal dysfunction. Meanwhile, the Class-2 group contained two genes whose manipulation worsened the phenotype, two genes that induced lethality, and 10 genes whose loss-of-function reduced neuronal dysfunction. Within the Class-2 group, we highlight that *Ptpro* (Ptp10D) was a particularly high-confident experimental candidate, with multiple loss-of-function constructs improving the neuronal health of the animals (**Figure 4b**).

These results suggest genes within the Class-1 and Class-2 groups may have the capacity to modulate neuronal dysfunction induced by mHTT. However, the small numbers of modifiers in each class makes it unclear whether Class-1 or Class-2 is more likely to contain genes that promote or counteract the homeostasis of neurons containing mHTT. Notably, the comparison of Class-1 and Class-2 genes with RNAi modifiers of neuronal dysfunction in transgenic *C. elegans* nematodes expressing an expanded (128Q) N-terminal exon-1 fragment of human HTT^32^ faced the same limitation and did not provide additional insight into this question; 399 genes ameliorated the phenotype and 263 genes worsened neuronal dysfunction. These data suggest that larger screens may be necessary to elucidate if there are differences in the number of pathogenic or compensatory changes in each class.

### The Class-2 gene *Atad-5* is connected to PCNA complex members deregulated in the direct striatal projection neurons of *post-mortem* human HD brains

To assess the human HD relevance of the Class-2 and Class-1 groups, we tested for overlaps with the lists of genes previously reported to be deregulated in the dSPNs of *postmortem* human HD brains either via direct analysis of dSPNs^2^ or single-cell sequencing methods ^23^. We detected no significant overlap, suggesting that Class-1 and Class-2 genes may be not relevant to gene deregulation in human HD dSPNs. However, manually adding high-confidence first neighbors for neighbor gene zipper analysis (see Methods/) highlighted a mouse-human short path of strong interest, where interconnected genes all primarily function into the PCNA complex, a protein complex that is key to the regulation of DNA repair in the course of DNA replication and that is strongly associated to somatic expansion in HD^33^ (**Figure 5**). This network cluster connects the Class-2 gene *Atad-5*, a PCNA unloader that is a modifier of somatic expansion in human HD blood samples^34^ to 6 human genes that are upregulated in the dSPNs of *postmortem* human HD brains^2^ including the regulators of nucleotide excision repair PCNA-interactor *CDKN1A* and histone acetyltransferase *EP300*^35^, the replisome components *DONSON* and *PRIM1*, the promoter of PCNA polyubiquitination *SHPRH* and the DNA polymerase *POLD1* (**Figure 5**). Additionally, this network cluster connects *Atad-5* to two genes of interest that in striatal neurons of *Hdh* mice shows upregulation at 6 month of age followed by reduction of this increase at 10 months of age^7^ including (*i*) *CTF18*, a clamp loader bound to PCNA^36^ that promotes touch receptor neuron dysfunction in human exon-1 *HTT C. elegans* nematodes^32^ and for which upregulation in the *Drd1* SNs of *Hdh* mice is pathogenic as indicated by *Geomic* analysis^7^, and (*ii*) *Brd4*, a negative regulator of PCNA unloading^37^ which binds and inhibits the ATAD-5 complex^38^ and is however not referenced in the STRING database. Given the knowledge on how ATAD-5, CTF18 and BRD4 modulate the activity of the PCNA complex and functional association of the PCNA complex with genetic modifiers of somatic expansion in HD such as *FAN-1*^39^, these results suggest that Class-2 genes may contain HD targets of strong interest for early intervention, *e.g*. blocking *CTF18* or modulating *Atad-5* expression might protect against HD via limiting the rate of somatic expansion in dSPNs.

**Figure 5.**
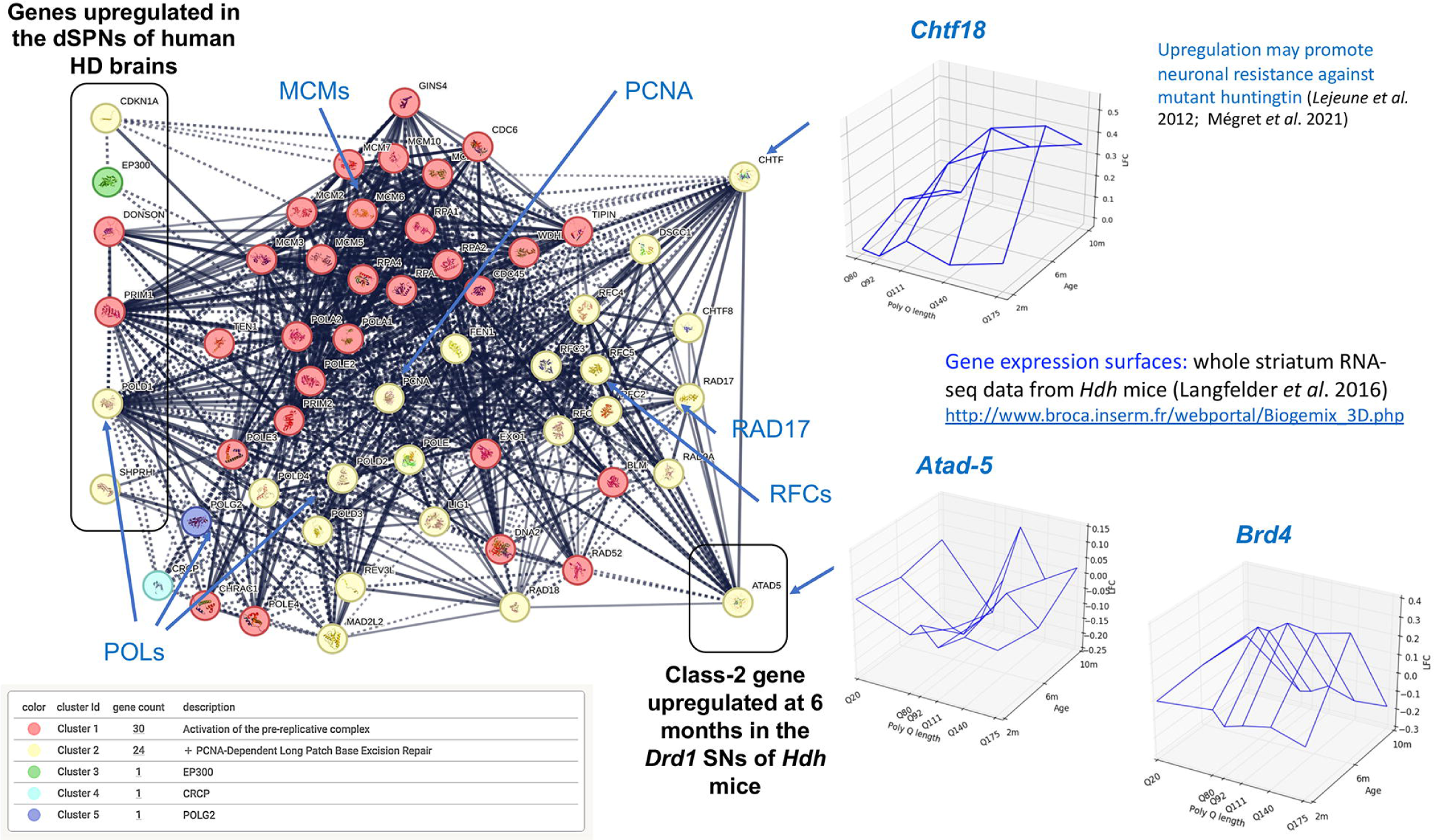
Behavioral screen of evolutionary-conserved Class-1 and Class-2 genes in transgenic HTT flies. **(a)** Selected genes from the Class-1 and Class-2 groups identified in mice were evaluated for their potential neuroprotective effects in *Drosophila* models of HTT. Fruit fly homologs for each group were identified, and selected genes were targeted for manipulation in *Drosophila*. Movement metrics such as speed of the flies over time were used as a behavioral (locomotor) assay as a readout of neuronal function. This approach aimed to determine whether alterations in the expression of specific Class-1 and Class-2 genes led to an improvement or decline in the progressive HTT-related phenotype. (**b)** Partial loss of function of Ptp10D, an ortholog of mouse gene Ptpro (Class-2 gene), improves the neuronal dysfunction phenotype in flies expressing HTT. Genetic perturbations shown are BDSC 53457 (left) and BDSC 5810 (right).

### *CXXC4* expression reduces p16INK4a levels in human HD and control iPS-cell-derived medium-spiny neurons

Our network model highlights the pioneer transcription factor *Sox11* (top CS-2 score) and the epigenetic regulator *Cxxc4,* also named *IDAX* (high CS-2 score in the barycenter area of temporal dynamics) as top genes for investigation of their protective effects and therapeutic potential against the dysfunction of striatal neurons in HD (**Figure 3c**). Since *Cxxc4* may promote synaptic activity via repression of TET2 and DNA methylation^29^, and since repression of TET2 may promote synaptic activity^40^ and protect from neurodegenerative diseases such as Parkinson’s disease ^41,42^ and neurodevelopmental disorders such as cerebral palsy^43^, we tested whether *CXXC4* expression may counter the detrimental effect of HD mutations on neuronal homeostasis and activity of human medium-spiny neurons (MSNs). To this end, we tested the effect of lentiviral overexpression of human *CXXC4* on the levels of p16INK4a in human isogenic HD iPS cell-derived MSNs, namely human HD72Q iPS cell-derived MSNs and their CAG-corrected counterpart C116 MSNs, a model that show increase of p16INK4a levels in HD72Q MSNs^11^. The expression of *CXXC4* reduced the abnormal levels of p16INK4a in human HD72Q iPS cell-derived MSNs and the basal levels of p16INK4a in C116-derived MSNs (**Figure 6a, Supplementary Figure 2a-b**), an effect exemplified in unrelated (non-isogenic) HD (HD-60Q) and control (18Q) lines (**Supplementary Figure 2c**), suggesting that *CXXC4* expression may promote MSN health.

**Figure 6.**
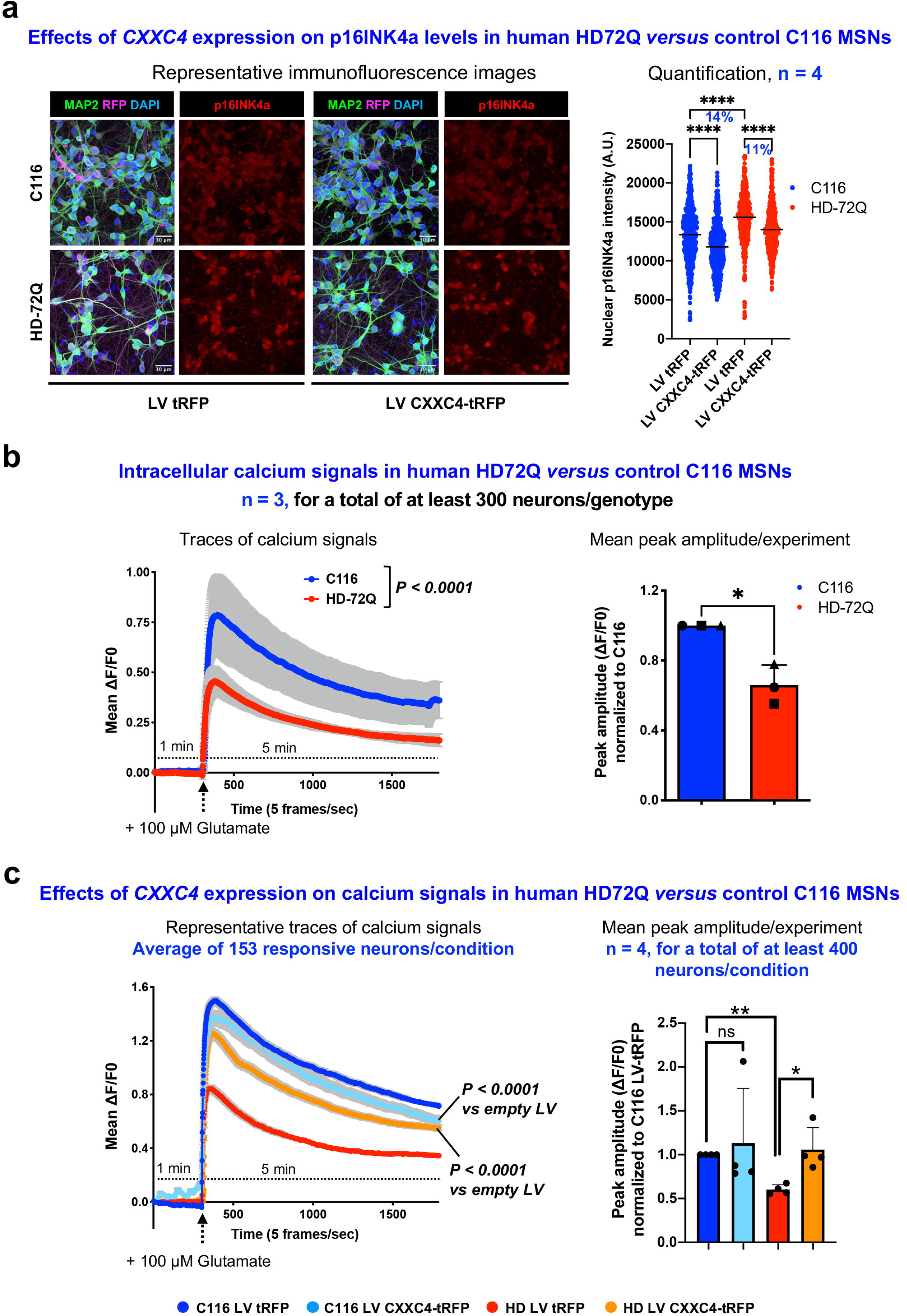
Effects of *CXXC4* expression on the cell senescence features and response to glutamate stimulation of isogenic human iPS cell-derived medium spiny neurons. (**a**) Left panels show representative immunofluorescence images of human HD72Q and control C116 iPS cell-derived MSNs transduced with LV CXXC4-tRFP or control LV tRFP and stained for MAP2, RFP and the senescence marker p16INK4a. Right panel shows quantification of nuclear p16INK4a intensity per cell (C116 LV tRFP: n = 667 cells, C116 LV CXXC4-tRFP: n = 631 cells, HD LV tRFP: n = 604 cells HD LV CXXC4-tRFP: n = 617 cells). *CXXC4* expression reduces nuclear p16INK4a levels in both HD and C116 neurons. Paired *t-*test, \*\*\*\**P* < 0.0001. 1. (b) Left panel shows intracellular calcium signals (mean ΔF/F ) recorded in responsive human iPS cell-derived MSNs following glutamate stimulation. Intracellular calcium signals to glutamate are significantly reduced in HD72Q compared to control C116 MSNs. Data are mean±SEM. Two-way ANOVA followed by Sidak’s multiple comparisons, *P <* 0.0001. Right panel shows quantification of mean peak amplitudes normalized to control cells C116. HD neurons show a 35% reduction in mean peak amplitude compared to C116 neurons. Data are mean±SD. Paired *t-*test, \**P* < 0.05. (**c**) Left panel shows representative traces of intracellular calcium signals (mean ΔF/F of responsive neurons) following glutamate stimulation as recorded in human iPS cell-derived MSNs transduced with either LV CXXC4-tRFP or control LV tRFP. Lentiviral expression of *CXXC4* strongly re-instates the response to glutamate in HD MSNs with no significant change detected for the response to glutamate in C116 MSNs. Data are mean±SD. Two-way ANOVA followed by Sidak’s multiple comparisons, \*\*\*\**P <* 0.0001 versus untreated. Right panel shows quantification of mean peak amplitudes normalized to C116 LV tRFP. In HD neurons, *CXXC4* expression increases peak amplitudes to C116 levels. Data are mean±SD. Paired *t-*test, \**P* < 0.05, \*\**P* < 0.01, ns: not significant.

### *CXXC4* expression restores the response of human HD iPS-cell derived mediumspiny neurons to glutamate stimulation

Next, we tested the effect of *CXXC4* expression on the response of human iPS-cell derived MSNs to glutamate stimulation using live imaging of intracellular calcium waves. Human HD72Q iPS cell-derived MSNs show defective response to glutamate stimulation compared to C116-derived MSNs (**Figure 6b, Supplementary Figure 2a-b**). The expression of *CXXC4* strongly restored glutamate excitability of human HD72Q iPS cell-derived MSNs with no significant change detected in C116 MSNs (**Figure 6c, Supplementary Figure 2a-b**). Together with the results on p16INK4a levels (**Figure 6a**), these results provide a proof-of-concept for the therapeutic potential of *CXXC4* expression and its capacity to restore the homeostasis and function of human MSNs in HD.

## DISCUSSION

Studies in *Hdh* mice suggest that HD mutations may translate into accelerated aging of striatal neurons on molecular levels such as epigenetic levels^8^, prematurely altering the expression of a large number of genes in the same direction as aging does, which may notably apply to gene downregulation. However, the temporal dynamics of this phenomenon has remained elusive in specific striatal neurons populations such as *Drd1* SNs. Our TAGL and TAGI data indicate that while aging-like transcriptional downregulation may be significant phenomenon in the *Drd1* SNs of *Hdh* mice, this phenomenon may be naturally mitigated as these mice progress from weakly to strongly symptomatic phases of the disease.

The mechanisms that may underlie the molecular changes that in striatal neurons are inverse compared to those in aging, and their biological significance, also remains elusive. This question noticeably applies to understanding the impact on the HD process of genes that are affected by aging and that are actually upregulated in the context of HD. Our TAGL and TAGI data reveal that transcriptional aging inversion effects of HD do not involve age-related genes at large, but may develop as a set of discrete and selective changes in the *Drd1* SNs of *Hdh* mice where a significant (50%) proportion of these changes are maintained as mice become increasingly symptomatic and may better explain disease progression compared to the dynamics of transcriptional-aging like effects.

Although molecular aging acceleration on transcriptional and epigenetic levels is an emerging feature of HD in *Hdh* mice^8^, it remains unclear whether striatal neurons in the human HD brain may undergo a similar process. Additionally, while cellular compensation is a universal and first-line response to aging and disease-related insults that makes a large use of cellular longevity mechanisms to promote cellular resilience, whether striatal neurons in the human HD brain may develop compensatory mechanisms that involve age-related genes is unknown. While molecular aging in brain cells may fundamentally differ between the mouse and human brain in terms of spread, dynamics and variability, our TAGL and TAGI data compared to *post-mortem* human HD brain data for MSNs suggest that aging like and inversion effects may be relevant to the molecular pathology of HD in human brains, raising the possibility that aging like and aging inversion may be at work in the human HD brains in a discrete, localized yet biologically important manner. Our data on the members of the PCNA complex also raise the possibility that the cellular mechanisms altered in *post-mortem* human HD striatal neurons can also be those that are altered by normal aging and by HD in mouse striatal neurons, and that some of these pathways may lie at the heart of the HD process as they relate to the regulation of somatic expansion, as suggested herein. However, the systematic investigation of these features remains limited by the lack of age-dependent gene expression data in striatal neurons isolated from *post-mortem* human brains.

3’UTR accumulation has been associated to aging of mouse striatal neurons^17^. Our data reveal that ribosomal mRNAs downregulated in these neurons are more likely affected by 3’UTR accumulation, highlighting a class of genes that may be strongly affected by aging in *drd1* SNs. Moreover, our data reveal that ribosomal mRNAs deregulated by HD in mouse *Drd1* SNs are more likely to be not only those that are deregulated in the course of normal aging but also those are affected by 3’UTR accumulation in normal aging. This rule noticeably applies to 83 genes that are affected by 3’UTR accumulation in aging, down-regulated in aging as well as *Hdh* mice, and associated with dendritic spine formation. These observations suggests that common events underlie aging and HD not only on transcriptional levels but also on post-transcriptional levels. This possibility will be further documented as 3’UTR accumulation data from *Drd1*-SNs of *Hdh* mice may become available. Interestingly, mapping 3’UTR accumulation data associated with normal aging onto networks associated with gene dysregulation in *Drd1* expressing striatal neurons of *Hdh* mice highlighted two classes of genes (Class-1 and Class-2 genes) that are interconnected and which size as well as functional annotations are CAG repeat length-dependent. Interestingly, Class-1 Q111 and Class-1 Q(170-175) genes partially overlap, a pattern that also applies to Class-2 genes. While Class-1 genes for Q111 are enriched in genes that protect from and genes that promote striatal neuron death in *Hdh* mice^22^, Class-1 genes for Q(170-175) are enriched only in genes that promote striatal neuron death, suggesting that Class-1 genes may shift from promoting a mix of pathogenic and compensatory effects to promoting direct pathogenicity as a striatal neuron faces increasingly large CAG repeats. Class-2 genes for Q111 are also enriched in genes that protect from and genes that promote striatal neuron death in *Hdh* mice, but the lack of functional shRNA screen data obtained in *Hdh* mice for these genes prevent to determine whether Class-2 Q(170-175) genes may show a CAG repeat-dependent shift to promote compensation. Interestingly, Class-2 Q(170-175) genes are associated with cell homeostasis (as inferred from annotations), and they contain genes of strong interest such as *CXXC4*, an epigenetic regulator that may repress TET2^29^, shown herein to protect human iPS cell-derived SNs against senescence-like phenotypes and to re-instate a normal response to glutamate stimulation in human HD iPS cell-derived SNs. Would restoring the response to glutamate stimulation be problematic in the context of the glutamate excitotoxicity hypothesis in HD^44^? The role of glutamate excitotoxicity in HD remains debated as this phenomenon — which is compensated by astrocytic uptake via GLT1 and GLAST— has been primarily studied *in vitro* using synaptosomes and may not be relevant in vivo^44^. The expression of *CXXC4* may therefore provide an efficient approach to sustain the MSN resiliency capacity via epigenomic remodelling and promotion of synaptic transmission, notably in the early or presymptomatic phases of the disease, and notably in MSNs that do not carry very large, somatically-expanded CAG repeats^2,23^, *i.e*. CAG repeats that could disrupt the capacity of the striatum to resist HD. Since a mechanism of action of CXXC4 is the repression of TET2^29^, and since inhibition of neuronal TET2 may enhance hippocampus-dependent memory^40^ and protect against neurodegenerative diseases such as Parkinson’s disease^41,42^, our data identify treatment with small compound molecules such as TET2 inhibitors as a strategy to restore MSN resilience capacity against HD. Additionally, our data suggest that expressing *CXXC4* might have therapeutic potential to treat neurons affected by neurodegeneratives diseases such as Parkinson’s disease, Alzheimer’s disease and repeat expansion disorders such as spinocerebellar ataxias. Finally, our data on CXXC4 protection of HD-vulnerable neurons such as striatal neurons add to the proposed links between repeat expansion and DNA hypermethylation at repeat loci^45^ and between DNA methylation levels and molecular pathogenesis of HD^46^.

In summary, our data reveal that, the transcriptional effects of HD in mouse *Drd1*-expressing striatal neurons do not involve age-related genes at large. Except perhaps to aging-like gene downregulation, the effect of HD on age-related genes mostly functions in the form of discrete and highly-dynamic patterns that involve specific groups of age-related genes, whether it may be for driving or resisting the HD process. Our data also reveal that genes that are down-regulated or affected by 3’UTR accumulation in aging, but that are upregulated in HD-vulnerable neurons such as *Drd1*-expressing striatal neurons may provide a source of new targets for early intervention in HD, notably to restore the capacity of MSNs to resist HD, as shown for *CXXC4*.

## METHODS

### Source data

We analyzed TRAP-seq data (ribosomal mRNA data) obtained from the *Drd1* SNs of the allelic series of HD knock-in (*Hdh*) mice covering Q20, Q111, Q170 and Q175 at 6 months of age, with 10 replicates per disease condition^15^ and ribosomal mRNA data obtained in the *Drd1* SNs of wild-type mice at postnatal day 42 (PN42) and at 24 months of age^17^. Both sets of read-count data were analyzed using the R package DESeq2 (version DESeq2_1.36.0) to generate the corresponding log-fold-change (LFC) values. For data from *Hdh* mice, Q20 data were used as a reference, whereas for wild-type aging, PN42 data were used as a reference. We also used 3’UTR accumulation RNA-seq data^17^ to annotate the resulting signatures.

### Curation of prior knowledge relevant to aging and stress resilience for enhanced biological annotations of gene sets

To complement information on biological content that can be retrieved in databases such as Enrichr^25^ or stringDB^47^ and with the aim to test for disease relevance, we compiled 30 datasets from publicly available curated databases and previously published studies related to neurodegenerative disease pathogenesis, aging, and stress resilience. The collection included functional screens in mouse models of HD^22^ and the integration of gene expression with functional screening data in *Hdh* mice^7^. FOXO3 targets were included from normal mice^19^ and from human HD neural stem cells^11^. We also considered *Htt* partner proteins^18^ and a striatal HD gene signature of 266 genes assembled by the CHDI foundation^48^ (https://www.hdinhd.org/). We further incorporated 479 genes identified as modifiers of somatic *mHtt* CAG-expansion rates in Q140 mice (FDR < 0.1), which showed at least a 20% change in expression^21^. Genes associated with long CAG repeat expansions (>150 units) were also included, namely phase C+ (upregulated), phase C– (downregulated), and phase D (de-repressed) genes^23^. In addition, we integrated genes reported as dysregulated in iMSN and dMSN of the cortex of human HD donors^2^. Finally, we included gene co-expression modules previously identified by weighted gene co-expression network analysis (WGCNA) of whole-striatum RNA-seq data across CAG repeat lengths and age points in *Hdh* mice^16^. Modules significantly associated with CAG length (|Z| > 5) were retained. Downregulated eigengenes were observed in modules M2, M25, M11, M34, and M52, whereas upregulated eigengenes were observed in modules M20, M9, M43, M7, M10, M39, M1, and M46.

The significance of the overlaps was calculated using Fisher’s exact test (scipy.stats.fisher_test, scipy version 1.16.2) using the combination of the all the dysregulated genes in HD and in aging as a universe for the TAGl/TAGi effect enrichment analysis. In the case of Class-1 and Class-2 genes, we considered the filtered MouseNet v2 network as explained in Section 3.4 as the full background for the significance analyses. Obtained p-values were adjusted for multiple testing for an alpha level of α = 0.05 (Benjamini-Hochberg correction).

### Identification of temporal dynamics of gene dysregulation in *Drd1*-expressing striatal neurons of *Hdh* mice

#### Retrieval of temporal information

To infer the temporal dynamics of gene dysregulation in *Drd1*-expressing striatal neurons, we improved our previously described algorithm *Geomic* (Mégret et al., 2021) to version 1.1. *Geomic* performs shape deformation analysis to align CAG repeat- and age-dependent whole-striatum dysregulation curves ^16^ with cell-type–specific dysregulation curves derived from *Hdh* mice data ^15^. Starting from cell-type–specific data are typically available for a single age point, *Geomic* v1.1 infers the most likely temporal dynamics by modeling the distribution of deformation distances between whole-striatum and cell-type specific curves as a mixture of three Gaussian components. These components represent genes where the cell type contributes strongly (short distance), partially (medium distance), or minimally (long distance) to the whole-striatum signal. We fitted this mixture model using the GaussianMixture function from *sklearn.mixture* (n = 3) and confirmed the presence of three distance classes based on BIC and AIC criteria. Distances between bulk and cell-type curves were computed with the Geomic suite^7^. Among 4,300 genes significantly dysregulated for at least two CAG repeat lengths at six months and showing a cell-type-specific expression, *Geomic* v1.1 successfully assigned 3,844 whole-striatum expression surfaces to at least one cell type, including 2,999 assignments to *Drd1* SNs. Each gene has up to four distance values (one per cell type). For each distance, we computed the probability of belonging to the short-, medium-, or long-distance distribution. Genes were assigned to a cell type when the probability of belonging to the short-distance population exceeded 0.7, corresponding to curves that are very close or nearly parallel between bulk and cell-type data. This procedure yielded one, several (up to four), or no cell-type assignments per gene, depending on the posterior probabilities.

#### Identification of temporal dynamics

To determine whether the genes we identified in the different subsets (TAGL, TAGI, Class-1, and Class-2) are maintained or reduced over time in the *Drd1*-SNs of *Hdh* mice, we first calculated the differences between 2-month and 6-month LFC values (ΔLFC_6M-2M_ = LFC_6M_ -LFC_2M_) and between 6-month and 10-month LFC values (ΔLFC_10M-6M_ = LFC_10M_ - LFC_6M_) for the whole striatum data of genes assigned to at least *Drd*1-SNs^16^. The product of the signs of these two delta values define two categories where a positive product identifies a maintained response (either increasingly or decreasingly monotonic) whereas a negative product identifies an inversion of the temporal profile.

### Network analysis of bootstrapped RNA-seq data using spectral decomposition of the signal (Rando-SDS analysis)

We combined bootstrapping approaches with network analysis techniques, namely spectral decomposition of the signal, developing a method herein named Rando-SDS. Firstly, we generated 100 bootstrap replicates of the mRNA read-count data from *Hdh* mice via sampling with replacement, thus obtaining 100 simulated experiments per CAG-repeat, each containing 10 mice replicates like in the original experiments. Bootstrapping was repeated for all the CAG-repeat lengths covered in the allelic series of HD knock-in mice. For each simulated experiment, read-count data were analyzed using DESeq2 to generate the corresponding LFC values, with Q20 data as a reference. This analysis was repeated for all CAG-repeat lengths, resulting in 100 LFC-value files per CAG-repeat length. Network analysis based on spectral decomposition of the signal (SDS)^49^ was then applied to the LFC files obtained for the bootstrap replicates, using a code that we previously reported^14^ as part of the BioGemix framework that we developed for post-omics machine learning (see http://www.broca.inserm.fr/BrainC_database/). Briefly, SDS enables LFC values to be smoothed against node connectivity in reference probabilistic function networks and noise to be removed while false negatives are retained, reducing the complexity of RNA-seq data while boosting the biological precision of resulting models^32,50^. SDS analysis was performed against the probabilistic functional network MouseNet v2^51^. To ensure biological robustness, we only retained MouseNet v2 modules deemed to be robust enough (all except MM-GN, MM-PG and SC-CC). After performing SDS analysis, we obtained a dysregulated network per bootstrap replicate of the initial experiment and per CAG-repeat length, generating a total of 400 SDS networks (100 networks per CAG repeat length) in addition to the four original non-bootstrapped networks. To ensure the statistical robustness of SDS networks, we retained edges that are present in 90 out of the 100 SDS networks built from the simulated experiments (boot-strapped datasets). This filtering procedure was performed for the four CAG-repeat lengths, resulting in networks with a size that is CAG-repeat length dependent (Q50: 56 nodes, 35 edges; Q111: 473 nodes, 461 edges; Q170: 915 nodes, 1300 edges. Q175: 990 nodes, 1516 edges). Given that Q170 and Q175 correspond to similar CAG repeat lengths, and given that 70% of the Q170 network is contained in the Q175 network, we merged these two networks into a single network referred as to the Q(170-175) network.

### Mapping aging-related 3’UTR accumulation data to Rando-SDS networks

To map aging-related 3’UTR accumulation data on Rando-SDS networks of gene dysregulation in the *Drd1* SNs of H*dh* mice, we used information on 3’UTR accumulation in the *Drd1* SNs of 24 month-old wild-type mice^17^. Sudmant *et al.* found a significant ribosome-associated accumulation of the average read coverage in the non-coding 3’UTR region of 5300 genes in *Drd1* SNs of 24-month old mice, a phenomenon not detected in young mice neurons. Sudmant *et al.* quantified this accumulation using the termination codon ratio (R_tc_). Here, we curated a sublist of genes only with positive R_tc_ values (negative R_tc_ values represent no accumulation of 3’UTRs and were therefore not considered). The termination codon ratio information (a.k.a. 3’UTR accumulation) was used to build a model that distinguishes genes that are affected by 3’UTR accumulation in aging and downregulated in *Hdh* mice (*i.e*., Class-1 genes: R_tc_ > 0 and HD LFC < 0) from genes that present 3’UTR accumulation in aging but are upregulated in *Hdh* mice (*i.e*., Class-2 genes: R_tc_ > 0 and HD LFC > 0). To integrate the information on the strength of 3’UTR accumulation in aged mice and that on the strength of gene dysregulation in *Hdh* mice, we calculated two comparison scores, namely CS-1 score for Class-1 genes and the CS-2 score for Class-2 genes.

Although both HD LFC data and R_tc_ data follow gaussian distributions, their value ranges greatly differ from each other. Hence, data required normalization so that the magnitude of LFC and R_tc_ values for a particular gene were in an equivalent scale for the calculation of the CS-1 and CS2-scores. To this end, we tested several data standardization and normalization algorithms, which led us to opt for the quantile standardisation scaling that standardizes data by transforming it to follow a uniform distribution (QuantileTransformed method from python’s sklearn.preprocessing package) as quantile standardization reduces the impact of marginal outliers, herein providing a robust pre-processing method.

The CS-1 and CS-2 scores are calculated as the product of a ratio and a sigmoid function that attenuates undesired artifacts hampening the actual effect of the ratio (e.g. large scores caused by very small values in the denominator). The CS-1 score is the ratio between the HD LFC value and the R_tc_ value of a given gene multiplied by a R_tc_-dependent sigmoid (*i.e*., 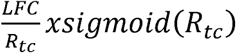), allowing to quantify the strength of Class-1 effects entailed by mutant huntingtin. The CS-2 score is the ratio of the R_tc_ value to the LFC value of a gene multiplied by a LFC-dependent sigmoid 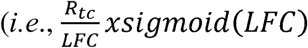, allowing to quantify the strength of Class-2 effects. The CS-1 and CS-2 scores were calculated for each CAG repeat length. The CS-1 and CS-2 scores for Q170 and Q175 were merged into a combined Q(170-175) score, which corresponds to the average of the Q170 and Q175 scores.

### Neighbor gene zipper analysis

To assess the human HD relevance of Class-1 and Class-2 genes (see above) gene lists associated with transcriptional changes in the *Drd1* SNs of *Hdh mice*, we tested whether the human orthologs of the Class-1 and Class-2 genes might be linked to genes deregulated in the dSPNs of *post-mortem* human HD brains^2^ via neighbor genes. To this end, we used high-confidence (TM+Experiments+databases+Expression data, score = 0.7) neighbors on the 1st shell as extracted from the STRING database^47^. We then manually screened the resulting paths for genes that, on both the mouse and human sides, may be associated with recently-discovered HD mechanisms. Paths of strong interest were further investigated at the protein complex level using neighbor gene zipper analysis in which we added 50 high-confidence 1st neighbors distributed on the 1^st^ shell or 2^nd^ shell in protein complexes —for any one Class-1 or Class-2 gene of interest— as extracted from the STRING database^47^ and assuming that a stacked protein complex may involve at least 50 proteins.

### Behavioral screen of top evolutionary-conserved Class-1 and Class-2 genes in transgenic HTT flies

To assess whether manipulation of the genes within the Class-1 and Class-2 signatures impact HD pathology, we utilized the well-established *Drosophila melanogaster* model of HD. Transgenic flies that express mutant N-terminal human HTT (HTTNT231Q128) pan-neuronally using the elav>GAL4 driver demonstrate deficits in neuronal function that that progress with age, which can be measured longitudinally using negative geotaxis assays^30,31^. After ranking the genes as previously described, we selected top genes from each category for *in vivo* screening. We identified *Drosophila* orthologs of the Class-1 and Class-2 genes using Drosophila RNAi Screening Center Integrative Ortholog Prediction Tool (DIOPT) version 9, filtering for the best matching candidates and those with a DIOPT score greater than or equal to 5. In total, we selected 29 orthologs from the Class-1 group and 23 orthologs from the Class-2 group for experimentation. Genetic constructs and alleles for the experiments were obtained from the Bloomington Drosophila Stock Center, totaling 90 constructs across the 52 genes.

To longitudinally assess neuronal function of the animals, we utilized a high-throughput robotic negative geotaxis assay, which has been previously described (Onur *et al.,* 2021). Briefly, four replicates of ten newly eclosed female flies per genotype were collected for experimentation and raised at 28°C. Flies were transferred to fresh food and underwent neuronal function screens every other day until day 15 post-eclosion. The climbing speed of the animals were graphed as a function of age. The climbing speed of each experimental genotype was compared to a set of healthy negative controls (elav>GAL4 crossed with the w1118 background or a non-targeting shRNA background) and diseased positive controls (elav>GAL4; HTTNT231Q128 crossed with the w1118 background or a non-targeting shRNA background). Statistical significance in changes in climbing speed were determined using a non-linear mixed effects regression model and a post-hoc for pairwise comparisons.

### Cell culture

Human iPS cell-derived from an HD patient (female 20 years old: 72Q/19Q, HD72Q) and their CAG-corrected counterpart (21Q/19Q, C116)^52^ and lines derived from an unrelated HD patient (female 29 years old: 60Q, CS21iHD-60 clone n8) and control individual (male 76 years old, CS25iCTR-18 clone n2) obtained from the Cedar Sinai consortium were used. iPSCs were differentiated into neural stem cells (NSCs) using the EB formation method (STEMCELLS Technologies). The differentiation into MSNs was performed using a modified version of a previously published protocol^53^. Briefly, NSCs were cultured on hESC-qualified Matrigel® (Corning) in Neurobasl medium (Gibco, # 21103049) supplemented with 2 mM GlutaMAX (Gibco), 100 U/mL penicillin, 100 μg/mL streptomycin, 1% B27 (Gibco # 17504044), 25ng/ml bFGF (PeproTech, #100-18B) and 10ng/ml LIF (PeproTech, #300-05) and passaged once a week. For neuronal differentiation, cells were dissociated with StemProAccutase and plated at a density of 200,000 cells/cm² on Matrigel-coated culture plates. Cells were pre-patterned for 7 days in DMEM/F12 supplemented with 2 mM GlutaMAX, 100 U/mL penicillin, 100 μg/mL streptomycin, 1% B27 without vitamin A (Gibco, # 12587010), 0.2 µM LDN-193189 (Tebubio, # 3481062443), 1.5 µM endo-IWR-1 (Biogems, # 1128234), and 20 ng/mL Activin A (PeproTech, #120-14E). On day 8, cells were plated on 100 µg/ml poly-D-lysine (Sigma, #P7280) and 10 µg/ml laminin (Sigma, #L2020) in SCM1 medium [DMEM/F12 supplemented with 2mM GlutaMAX, 1% B27, 1.8 mM CaCl_2_, 2 µM PD033291 (Bio-Techne, #4786/10), 200 µM Ascorbic Acid (Sigma, # PHR1008-2G), 300 µM γ-Aminobutyric acid (Sigma, #A2129-10G), 5 µM LM22A4 (Tocris, #4607/5), 3µM CHIR99021 (Sigma, # SML1046-5MG), 300 µM GABA, 10 µM Forskolin (Sigma, # F6886-10MG), 10 µM DAPT, 10 ng/mL BDNF (PeproTech, #450-02)]. Half-medium changes were performed every two-three days. On day 16, a full medium change was performed to SCM2 medium [1:1 mixture of DMEM/F12 and Neurobasal Medium supplemented with 2mM GlutaMAX, 100 U/mL penicillin, 100 μg/mL streptomycin, 1% B27, 1% N2 (Gibco # 17502048), 1.8 mM CaCl_2_, 3µM CHIR99021, 2 µM PD033291, 200 µM Ascorbic Acid, 5 µM LM22A4 and 10 ng/mL BDNF]. Half-medium changes were performed every two-three days until day 30. Differentiation into MSNs was assessed by immunofluorescence using antibodies against MAP2 (1:1000, Sigma # M4403), GABA (1:250, Sigma #A0310) and DARPP32 (1:100, Santa Cruz Biotechnology #SC-11365). Lentiviral transduction was performed using lentiviral vectors produced by the MiRCen platform (CEA Paris-Saclay, France). Human *CXXC4* cDNA, tagged with a V5 epitope at the N-terminus, was sequence-optimized and synthesized by GeneArt (Thermo Fisher Scientific). The cDNA were then subcloned into a third-generation lentiviral backbone under the control of the PGK promoter, followed by an IRES-TagRFP cassette to allow fluorescent visualization of transgene expression (pLVi-PGK-TagRFP-WPRE and pLVi-PGK-V5.CXXC4-IRES-TagRFP). Lentiviral particles were added to the culture medium at a multiplicity of infection (MOI) of 10, on day 30 of differentiation. Twenty-four hours post-transduction, the medium was replaced with fresh SCM2 medium, and cells were maintained under standard differentiation conditions for an additional 72 hours. MSNs were thus transduced for a total of four days prior to downstream analyses. Transduction efficiency was assessed by immunofluorescence using an anti-RFP antibody (1:200, GeneTEX #GTX628545) and by qRT-PCR using the primers as described below.

### qRT-PCR analysis

Total RNA was extracted from cells using the NucleoSpin RNA kit (Macherey-Nagel) and subsequently treated with DNase using the DNA-free DNA Removal Kit (Ambion) according to the manufacturers’ protocols. Equal amounts of RNA (1 μg) were reverse-transcribed into cDNA using the RevertAid First Strand cDNA Synthesis Kit (Thermo Fisher Scientific, K1622), following the manufacturer’s instructions. The resulting cDNA was diluted and used as a template for quantitative real-time PCR (qRT-PCR) analysis. qRT-PCR was performed on a LightCycler 480 Real-Time PCR System (Roche) using GoTaq qPCR Master Mix (Promega, A6002). All qRT-PCR reactions were run in triplicate with the following primers: RFP_For AAGAATCAAGGAGGCCGACA and RFP_Rev CTTGTACAGCTCGTCCATGC; LV-CXXC4_For TCAGCGCCATTCCAGCTCTCG and LV-CXXC4_Rev GGCGATCTGGAAGGCAGAATC.

### Immunofluorescence analysis for quantification of p16INK4a levels

MSNs were differentiated in μ-Slide 8 Well high ibiTreat (ibidi, #80806) coated with poly-D-lysine and laminin as described above. Cells were fixed with 4% paraformaldehyde (ThermoFisher Scientific) and 4% sucrose for 20 min at room temperature (RT) and then washed twice with DPBS. Blocking and permeabilization were performed simultaneously using 3% bovine serum albumin (BSA) and 0.2% Triton X-100 (Sigma-Aldrich) in DPBS for 1.5 hours at RT. Cells were incubated overnight at 4 °C with the following primary antibodies diluted in 1% BSA in DPBS: anti-p16INK4a (1:200, Abcam # Ab108349), anti-MAP2 (1:1000, Sigma # M4403) and anti-RFP (1:200, GeneTEX# GTX628545). The next day, cells were washed three times with DPBS for 10 min and incubated for 2 hours at RT in the dark with the following secondary anti-bodies: Cy5 AffiniPure™ donkey anti-rabbit (1:400, Jackson ImmunoRes # 711-175-152), Cy3 AffiniPure™ Goat anti-mouse IgG2a (1:400, Jackson ImmunoRes #115-165-206), and Goat anti-Mouse IgG1 cross-adsorbed, Alexa Fluor™ 488 (1:500, Thermo Fisher, #A-21121). Nuclei were counterstained with DAPI (Sigma, #D9542). After three additional DPBS washes ibidi mounting medium (ibidi, #50001) was added to preserve fluorescence. Imaging was performed using a Leica SP5 Confocal Microscope (Leica Microsystems, Wetzlar, Germany), with identical acquisition settings for each condition to ensure data comparability. Images were analyzed using ImageJ software (https://imagej.nih.gov/ij/). First, DAPI images were were processed to generate a binary mask defining nuclear regions of interest (ROI). These masks were then sequentially applied to the MAP2 and to the RFP channels to exclude non-neuronal, undifferentiated or non-transduced cells. Finally, the selected ROIs were applied to the p16INK4a channel to quantify nuclear fluorescence intensity. Background fluorescence was calculated as the mean intensity of areas outside the ROIs and subtracted from the measured values.

### Calcium imaging and analysis

Human iPS cell-derived medium spiny neurons (MSNs) were differentiated in μ-Slide 8 Well high ibiTreat (ibidi, #80806) and transduced with lentiviral vectors for four days as described above. For calcium imaging, cells were incubated with the calcium-sensitive dye Fluo-4 AM (1µM, Thermo Fisher Scientific, #F14217) diluted in recording buffer (129mM NaCl, 4mM KCl, 1mM MgCl_2_, 2mM CaCl_2_, 10mM Glucose, 10mM Hepes) containing 0.02% Pluronic F-127 (Thermo Fisher Scientific, #P3000MP) for 30 minutes at 37 °C. After three gentle washes with pre-warmed recording buffer, the ibidi slides were transferred to microscope for imaging. Live-cell calcium recordings were performed using a Zeiss Axiobserver microscope equipped with a pE4000 illuminator (CoollEd) triggering an Orca Fusion BT CMOS camera (Hamamatsu) and controlled by the HCImage live software (Hamamatsu). MSNs were imaged at room temperature for seven minutes at a rate of five frames per second (2 100 frames in total) using a 20X objective at 0.2Hz (50ms exposure time). At the beginning of each recording session, an image was captured in the RFP channel to select cells transduced with the lentiviral constructs. After a one-minute of baseline recording period (300 frames), 100 μM glutamate (Sigma, #G1626) was applied directly to the imaging well, and fluorescence was recorded continuously for an additional five minutes (1 500 frames). Finally, 100 µM KCl was added to the cells at frame 1 800 as a positive control to assess neuronal viability and activity. Fluorescence was then recorded for an additional minute (up to frame 2 100). Fluorescence intensity over time was analyzed using NETCAL software (http://www.itsnetcal.com/). Briefly, regions of interest (ROIs) corresponding to individual cells were automatically detected from images, and the fluorescent traces for each cell (ROI) were extracted. Data were further processed to extract peak amplitudes, and the response kinetics were analyzed and clustered using a custom in-house Python pipeline (see Code availability). Calcium responses were expressed as changes in fluorescence relative to baseline (ΔF/F ). For each cell, the first four frames were discarded due to their high oscillation and the baseline fluorescence (F ) was defined as the average fluorescence intensity between frames 5 and 20. Then, for each frame, ΔF/F was calculated using the formula: (F − F )/F , where F represents the fluorescence intensity at that time point. Only cells with peak responses between frames 200 and 400 that exceeded twice the baseline standard deviation were retained for further analysis. To assess responses to glutamate (injected at frame 300) and KCl (injected at frame 1800), the following additional criteria were applied: the response at frame 1750 must show a decrease of at least 35% compared to the peak value identified in the previous filtering step, the response at the midpoint between the peak and frame 1750 must be at least 15% lower than the peak and the response had to remain non-negative from frame 1750 onward. Cells that did not meet these criteria, including those with a low response to glutamate (peak ΔF/F < 0.15), were excluded from the analysis. Additionally, wells in which fewer than 10% of cells responded to glutamate were also excluded. For each differentiation, at least two ibidi slides were used, with two wells per condition. Within each well, the responses of all eligible neurons were averaged, and the standard error (SD) was calculated. To account for inter-experimental variability, the average peak amplitude within each well was normalized to the corresponding average peak amplitude of control cells.

### Statistical analysis

All experiments were performed at least using three independent neuronal differentiation cultures, unless otherwise stated in the figure. Statistical analyses were performed using GraphPad Prism (version 9.0, GraphPad Software, La Jolla, CA, USA). Data are presented as mean±SD, unless otherwise specified, based on the number of independent experiments indicated in the figure. For comparisons between two groups, significance was determined using a paired Student’s *t*-test. For analysis of cellular senescence marker levels, significance was determined using a paired Student’s *t*-test. For analysis of calcium traces, a two-way ANOVA followed by Sidak’s multiple comparisons test was performed. For analysis of mean peak amplitude, significance was determined using a paired Student’s *t*-test. A *P* value < 0.05 was considered statistically significant.

## DATA AVAILABILITY

TRAP-seq data used herein were previously reported, including TRAP-seq data obtained in *Drd1*-expressing striatal neurons of 6-month-old HD knock-in mice^15^, data representing 3’UTR accumulation data in *Drd1-*expressing striatal neurons of 24-month-old wild type mice^17^ and RNA-seq data obtained in the whole striatum of *Hdh* mice^16^. The data generated herein can be visualized in the form of interactive 2D 3D graphs at http://www.broca.inserm.fr/Molaging/index.php.

## CODE AVAILABILITY

The source code for Rando-SDS analysis (Rando-SDS package v 1.0), written using python, R and bash is available at http://www.broca.inserm.fr/Molaging/index.php and https://github.com/Brain-C-IBPS/RandoSDS. The source code used to classify calcium wave imaging curves is available at http://www.broca.inserm.fr/Molaging/index.php.

## ACKNOWLEDGEMENTS

This work was supported by Sorbonne Université, CNRS and INSERM, Paris, France and by the CHDI Foundation (grant number A-14814), Princeton, USA and CNRS Innovation (prematuration program Changing-HD), France (to C.N.) and by grants from NIH (grant number U01 AG0722439) and the Huffington foundation (to J.B.).

## AUTHOR CONTRIBUTIONS

M.A.L. performed data mining and bioinformatics data analyzes, analyzed the data and wrote the manuscript. F.F. performed biological experiments in human cells, analyzed the data and wrote the manuscript. T.M.A. helped with biological experiments in human cells and analyzed the data.

M.M. performed *Drosophila* studies, analyzed the data and wrote the manuscript. C.M. helped with machine learning. H.T. helped with the analysis of calcium wave imaging data. J.A. and J.R. contributed essential prior-knowledge data on HD signatures. L.E. contributed essential reagents (isogenic human HD72Q and C116 iPS cell lines). E.B. helped with data analysis. J.M.P. helped with data calcium wave imaging data acquisition and analysis. J.B. contributed essential data (*Drosophila* data), analyzed the data and wrote the manuscript. C.N. conceived the study, analyzed the data and wrote the manuscript. L.M. designed machine learning, analyzed the data and wrote the manuscript. All authors read and approved the manuscript.

## SUPPLEMENTARY INFORMATION

### Supplementary Tables

**Supplementary Table 1.** Differential expression analysis of HD and normal-aging TRAP-seq data in mouse *Drd1*-expressing striatal neurons, gene set category (aging-like and aging-inversion effects), *Geomic* v 1.1 assignation, temporal profile, whole-striatum LFC changes between 2 months and 6 months, and whole-striatum LFC changes between 6 months and 10 months.

**Supplementary Table 2.** Probabilistic overlap analyzes between aging-like or aging-inversion effects in the *Drd1*-expressing striatal neurons of *Hdh* mice and datasets of interest.

**Supplementary Table 3.** Probabilistic overlap analyzes between deregulated genes (TRAP-seq data) and genes affected by 3’UTR accumulation in the *Drd1*-expressing striatal neurons of 24-month-old wild-type mice^17^.

**Supplementary Table 4.** Probabilistic overlap analyzes between genes deregulated in *Hdh* mice (TRAP-seq data)^15^, genes deregulated in aging (TRAP-seq data)^17^, and genes genes affected by 3’UTR accumulation in aging^17^, all datasets from the *Drd1*-expressing striatal neurons.

**Supplementary Table 5.** Connectivity matrices for Rando-SDS networks defining Class-1 and Class-2 genes for CAG repeat size Q50, Q111 and merged Q(170-175).

**Supplementary Table 6.** Information for Class-1 and Class-2 genes, including gene-expression levels at different CAG repeat lengths, R_tc_ values representing the 3’UTR accumulation, *Geomic* v 1.1 assignation, temporal profile, bulk striatum LFC changes between 2 and 6 months, bulk striatum LFC changes between 6 and 10 months, and CS-1 and CS-2 scores.

**Supplementary Table 7.** Class-1 and Class-2 genes for Q111 and Q(170–175), with and without inclusion of first-neighbour interactors.

**Supplementary Table 8.** Overlaps between Class-1 or Class-2 genes and genes in datasets of interest.

**Supplementary Table 9.** Functional annotations for Class-1 or Class-2 gene nodes and their neighbours as indicated by *Geomic* analysis^7^ and by functional shRNA screening in *Hdh* mice^22^.

**Supplementary Table 10.** Overview of *Drosophila* screen data, including ortholog mapping, genotype/allele information, observed phenotypes, and statistical significance of genotype, time, and type of effects.

### Supplementary figures

**Supplementary Figure 1 related to Figure 3.** (**a**) Machine learning pipeline for characterizing cumulative effect class (CEC) and non-cumulative effect class (non-CEC) signatures in the *Drd1* SNs of *Hdh* mice. The analysis pipeline takes advantage of two datasets, the first corresponding to RNA-seq data from the *Drd1* SNs from 6-month-old *Hdh* mice bearing 4 CAG-repeat lengths, and the second to 3’UTR accumulation data from the *Drd1* SNs of 24-month-old wild type mice. In a first step, using statistical and bionetwork-based approaches for each CAG-repeat length, the pipeline classifies genes retained by the model into two different groups: genes subjected to down regulation in HD and 3’UTR accumulation in normal aging and genes subjected to up regulation in HD and 3’UTR accumulation in normal aging. In a second step, applying an improved version (v. 1.1) of *Geomic* analysis^7^, the pipeline performs gene-by-gene inference of temporal dynamics of CEC and non-CEC effects by exploiting whole striatum RNA-seq data of *Hdh* mice mice at 2 months, 6 months and 10 months of age, assigning the temporal information that is available in the whole striatum data to CAG-repeat dependent data of *Drd1* SNs. (**b**) Interconnections between the Q111 Class-1 and Class-2 networks in the *Drd-1* SNs of *Hdh* mice at 6-months. Class-1 nodes are represented in dark red, Class-2 nodes in dark green, nodes downregulated in HD but not present in the Class 1 signature are light red, while downregulated in HD but not present in the Class 2 signature are light green. Node size is proportional to the CS1 and CS2 scores of Class-1 and Class-2 genes, respectively. Most remarkable EnrichR annotations are also indicated. (c) Temporal dynamics of Class-1 and Class-2 effects in the *Drd1* SNs of *Hdh*-Q111 mice. Out of 389 genes retained by our model in the combined Q(170-175) network (246 Class-1 genes and 143 Class-2 genes), 222 genes (168 Class-1 genes and 54 Class-2 genes) were assigned at least to the *Drd1* SNs. These genes are further classified into 4 temporal subclasses: reduced Class-1 effects, maintained Class-1 effects, reduced Class-2 effects and maintained Class-2 effects. Panels feature 2D representations of each of the four classes, where the x-axis corresponds to the difference between LFC values at and at 6 months compared to 2 months (ΔLFC_6M-2M_) and y-axis to the difference between LFC values 10 months compared to 6 months (ΔLFC_10M-6M_): (top left) Class-1 genes whose temporal profile is reduced across age points [39 genes]; (top right) Class-2 genes whose temporal profile is reduced across age points [3 genes]; (bottom left) Class-1 genes whose temporal profile is maintained age points [11 genes] and (bottom right) Class-2 genes whose temporal profile is maintained across age points [2 genes]. The straight line in each panel represents the y = x line where the two ΔLFC have the same value. The color gradient represents the combined Q(170-175) CS-1 or CS-2 score at 6 months of age. For each panel, a number of interesting genes have been highlighted.

**Supplementary Figure 2 related to Figure 6.** Quality control of MSN differentiation and lentiviral transduction. (**a**) Representative immunofluorescence images of human iPS cell-derived MSNs C116 (CAG-corrected: 21Q/19Q) and HD (72Q/19Q) stained for gamma-aminobutyric acid (GABA) and DARPP32. Scale bar: 30 µm. Right panel shows the percentage of cells positive for each marker. Data are mean±SD. (**b**) Upper panels show qPCR analysis of relative mRNA expression levels of *RFP* and *CXXC4* in transduced cells compared to untreated cells (UT). Lower panel shows the percentage of MAP2- and RFP-positive cells based on immunofluorescence analysis, indicating successful neuronal differentiation and effective transduction. Data are mean±SD. (**c**) Quantification of nuclear p16INK4a intensity per cell in unrelated HD-60Q and control-18Q iPS cell lines (18Qn2 LV tRFP: n = 184 cells, 18Qn2 LV CXXC4-tRFP: n = 126 cells, HD60Qn8 LV tRFP: n = 121 cells, HD60Qn8 LV CXXC4-tRFP: n = 59 cells). Experiment indicating that *CXXC4* expression may reduce nuclear p16INK4a levels in both human HD60Qn8 and control 18Qn2 iPS cell-derived MSNs. Paired *t-*test, \*\*\*\**P* < 0.0001.

