## Supplementary figures and images for "Modeling Huntington’s disease against age-related genes reveals *CXXC4* as an epigenetic target to restore DRD1 striatal neuron activity"

### Supplementary Fig 1

Supplementary Figure 1 related to Fig. 3

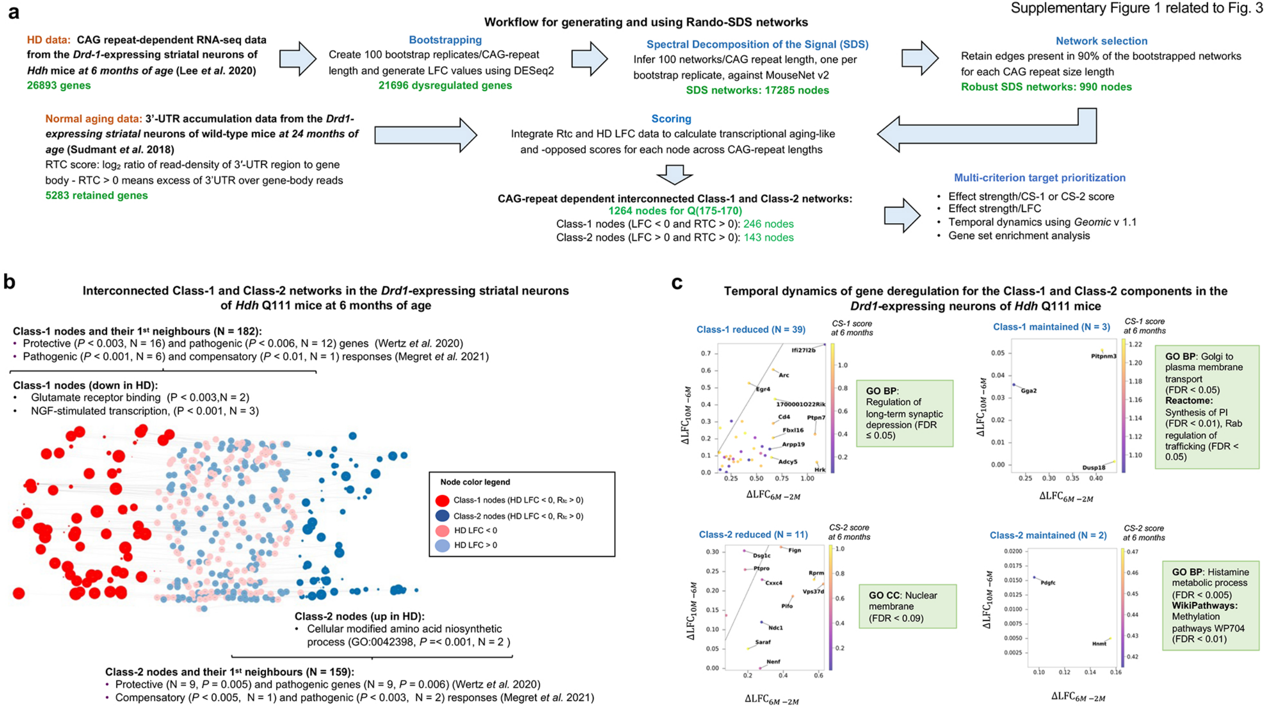
