## Supplementary Fig 2 for "Modeling Huntington’s disease against age-related genes reveals *CXXC4* as an epigenetic target to restore DRD1 striatal neuron activity"

**a**

### Human isogenic iPS cell-derived medium spiny neurons

**Differentiation quality control - Representative data (N = 3)**

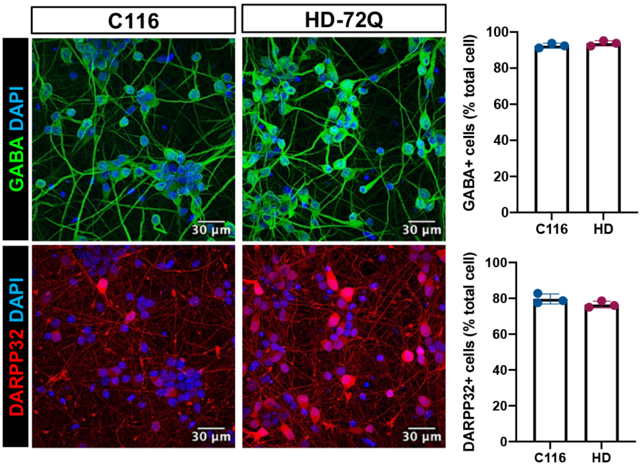


**b**

### Quality control for lentiviral transduction

qRT-PCR analysis

Immunofluorescence staining analysis

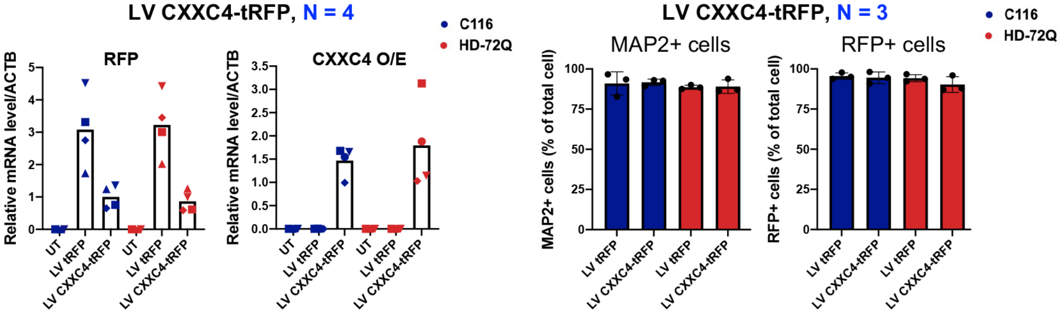


**c**

### Effects of *CXXC4* expression on p16INK4a levels in human HD 60Qn8 *versus* control 18Qn2 iPS cell-derived medium spiny neurons

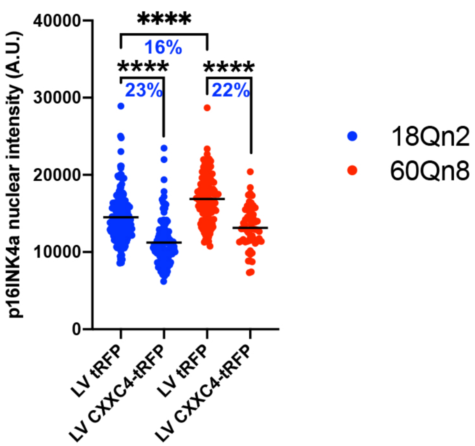
